# Cross-modal interactions at the audiovisual cocktail-party revealed by behavior, ERPs, and neural oscillations

**DOI:** 10.1101/2022.09.30.510236

**Authors:** Laura-Isabelle Klatt, Alexandra Begau, Daniel Schneider, Edmund Wascher, Stephan Getzmann

## Abstract

Theories of attention argue that objects are the units of attentional selection. In real-word environments such objects can contain visual and auditory features. To understand how mechanisms of selective attention operate in multisensory environments, we created an audiovisual cocktail-party situation, in which two speakers (left and right of fixation) simultaneously articulated brief numerals. In three separate blocks, informative auditory speech was presented (a) alone or paired with (b) congruent or (c) uninformative visual speech. In all blocks, subjects localized a pre-defined numeral. While audiovisual-congruent and uninformative speech improved response times and speed of information uptake according to diffusion modeling, an ERP analysis revealed that this did not coincide with enhanced attentional engagement. Yet, consistent with object-based attentional selection, the deployment of auditory spatial attention (N2ac) was accompanied by visuo-spatial attentional orienting (N2pc) irrespective of the informational content of visual speech. Notably, an N2pc component was absent in the auditory-only condition, demonstrating that a sound-induced shift of visuo-spatial attention relies on the availability of audio-visual features evolving coherently in time. Additional analyses revealed cross-modal interactions in working memory and modulations of cognitive control. The preregistered methods and hypotheses of this study can be found at https://osf.io/vh38g.

## 1. Introduction

Theories of attention argue that the units of attentional selection – both in vision and audition – are individuated perceptual objects (Shinn-Cunningham, 2008). In order to selectively attend to what is perceived as a coherent percept, the brain must first solve the problem of binding stimulus features that belong together, while separating them from potentially overlapping signals not coming from the same source. This computational problem is commonly referred to as auditory scene analysis in audition (Bregman, 1990) and as image segmentation in vision (Driver et al., 2001). Binding linked features from a common source does not only pose a problem within sensory modalities, but has to be extended to a multisensory dimension, particularly in natural environments, in which auditory and visual signals are seamlessly bound into a coherent audiovisual percept (Bizley et al., 2016). Consequentially, the demands for attentional selection are multimodal in nature. For instance, having a conversation at a crowded restaurant becomes easier when attending to your friend’s face and mouth movements in addition to their voice. In contrast, looking at somebody else’s face while eavesdropping on the conversation at the neighboring table will be more effortful (Bizley et al., 2016). These real-world examples demonstrate how multisensory objects might be more easily segregable from competing stimuli and how principles of object-based attention are straightforwardly applicable to situations requiring audiovisual processing (Lee et al., 2019). Yet, to date, the selection of task-relevant objects from a crowded scene and its neural underpinnings have been primarily investigated in a unisensory context (see e.g., for an overview Lee, 2017).

An object-based attention account makes one fundamental prediction: that is, when attending to one feature of an object, the unattended features of the same object will also be prioritized (Blaser et al., 2000; O’Craven et al., 1999). Applying the same logic to multisensory objects, this means that attention should also spread to simultaneous signals across modalities. In line with this prediction, in a combined EEG and fMRI study, Busse and colleagues (2005) showed enhanced activity related to the processing of centrally presented, irrelevant tones when the tones co-occured with a covertly attended, lateralized visual target stimulus compared to when paired with an unattended visual stimulus. Specifically, ERP subtractions isolated a multisensory attention effect for the tones over fronto-central areas, emerging at around ~220 ms and lasting until ~700 ms following stimulus onset, while fMRI results confirmed that the effect entailed enhanced activity in auditory sensory cortex. The authors interpret their results as a late, object-based attentional selection process, indicating that the attended visual as well as the initially unattended auditory stimulus at a discrepant location are grouped together. Along similar lines, a study by Van der Burg et al (2011) investigated ERP correlates of visuo-spatial attentional selection in an audiovisual search paradigm. They asked participants to monitor a dynamic search display of several diagonal lines that changed orientations at pseudo-random intervals. Subjects had to respond whenever a distractor line changed into a target line with either a horizontal or vertical orientation. A concurrently presented sound either occurred in synchrony with the target orientation change (AV-target) or simultaneously with a distractor orientation change (AV-distractor). Again, in line with an object-based attention account, a sound-elicited N2pc component – indicating the focusing of visuo-spatial attention (Luck & Hillyard, 1994) – occurred irrespective of whether the simultaneously presented visual stimulus was a target or an irrelevant distractor (Van der Burg et al., 2011). The authors interpret the presence of an N2pc in AV-target as well as AV-distractor trials as evidence for an automatic integration of the sound and the visual stimulus.

Notably, the studies described above all presented the sounds and the visual stimuli at disparate locations, thus creating illusionary percepts of a combined audiovisual stimulus emerging from the same spatial location. Moreover, both van der Burg et al (2011) and Busse et al. (2005) chose auditory input as the task-irrelevant modality, hence, leaving open to what extend the observed phenomena are bi-directional. Lastly, they used temporally coherent, but otherwise disparate cross-sensory features that have no presumed natural connection (e.g., checkerboard stimuli and a tone pip); hence, they disregard the possibility that both modalities could provide complementary or redundant information that is linked in a meaningful way.

Addressing those open questions and constraints, this preregistered study (https://osf.io/vh38g) aimed to shed light on the neural basis of attentional focusing within multisensory environments, using natural audiovisual speech stimuli. Across three different task-blocks, participants were asked to indicate the location of a pre-defined target numeral among a set of two simultaneously presented numerals (see Figure 1). Responses were given with their dominant hand. The acoustic speech stimuli were either presented (a) without any visual information (auditory only), (b) together with congruent visual information (i.e., a recording of a speaker’s face, in which the lip movement matched the presented auditory information) or (c) together with uninformative visual information (i.e., a recording of a speaker’s face, in which the lip movement did not match, but pronouncing a ‘ba’ lip movement). Noteworthy, as in previous investigations (Begau et al., 2021, 2022), the visual stimulus material consisted of natural recordings of a speaker’s face, allowing for an ecologically valid simulation of a multi-speaker scenario.

**Figure 1.**
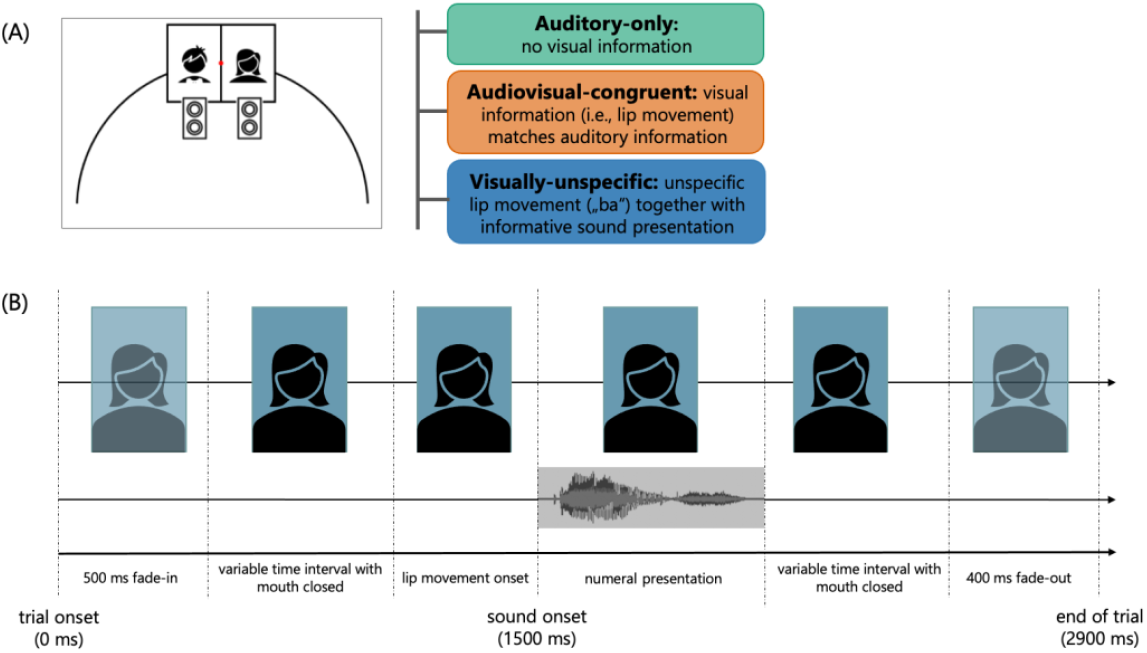
Experimental Design. (A) Overview of the experimental setup and conditions. For video presentation, two monitors were mounted to the right and left of central fixation at a viewing distance of 1.5 m. A loudspeaker was mounted underneath each monitor. Acoustic speech was either presented (i) without visual information, (ii) together with congruent visual information or (iii) together with uninformative visual information. Conditions were presented block-wise. In all blocks, participants were asked to indicate the location (left vs. right) of a pre-defined target numeral. Participants were instructed to respond as accurately and as fast as possible. Responses were given by pressing one out of two buttons with the index and middle finger of their dominant hand. (B) Exemplary illustration of a video sequence of a female speaker uttering short verbal stimuli. Please note that in each trial two stimuli were concurrently presented. The depicted sequence only shows an exemplary video sequency at one of the two locations. The speaker’s face was replaced by a placeholder object. For an image of the actual stimulus material, please see: https://osf.io/asx5v/.

This allowed us to answer three overarching research questions: First, we aimed to investigate to what extend the presence of congruent or uninformative visual information aids the listener to select the relevant sound from a mixture of competing inputs in a multi-talker environment. To this end, we examined behavioral performance measures as well as N2ac amplitudes as a correlate for auditory attentional focusing (Gamble & Luck, 2011). Second, the present task setup allowed us to investigate whether audiovisual events automatically induce both auditory and visual selective attentional orienting, even if only the auditory modality contains task-relevant information; alternatively, a-prior knowledge about the fact that visual speech content was uninformative (i.e., an unspecific ‘ba’ lip movement) could suppress visuo-spatial attentional orienting. To this end, in addition to the auditory N2ac component, the N2pc component serves as a correlate for visual attentional selection (Luck & Hillyard, 1994). Lastly, inspecting N2ac and N2pc latencies allows conclusions regarding the question whether the timing of auditory and visual attentional selection in audiovisual scenes is likewise dependent on the informational content (i.e., meaningfulness) of the visual input.

Taken together, to unravel the interplay and temporal dynamics of unisensory attention mechanisms in an audiovisual multi-talker environment, our pre-registered hypotheses focused on behavioral effects as well as the N2ac and N2pc components. However, additional exploratory analyses also considered subsequent, lateralized components to uncover potential cross-modal interactions in working memory, related to the in-depth analysis of task-relevant information (Jolicœur et al., 2008; Mazza et al., 2008; Töllner et al., 2013). Moreover, a growing body of work emphasizes the relevance of neural oscillations for multisensory processing (reviewed by Keil & Senkowski, 2018). In particular, modulations of theta power (4-7 Hz) have been associated with the detection of audiovisual incongruency and the need for cognitive control during audiovisual integration (Begau et al., 2022; Morís Fernández et al., 2018). Hence, in addition, time-frequency dynamics underlying attentional orienting in this multisensory cocktail-party scenario were explored.

## 2. Results

### 2.1 Concurrent visual speech facilitates sound localization, even if visual speech content is uninformative

If complementary visual cues facilitate target individuation and selection from a complex scene, this should be evident in terms of faster response times and an increased rate of information uptake (i.e., higher drift rates, a parameter estimated by diffusion modeling) for congruent audiovisual information (compared to auditory-only stimulus presentation). In contrast, with respect to accuracy, we expected participants to perform close to ceiling.

Figure illustrates the mean proportion of correct responses, mean response time, and mean drift rate per condition and for each individual participant. As expected, all participants performed very accurately. Accordingly, the analysis of accuracy revealed no significant differences between conditions, χ^2^(2) = 3.985, *p* = .136, W = 0.055. A complementary Bayesian analysis suggest no strong conclusions in favor for or against the null hypothesis, given that the Bayes Factor lays in the range of anecdotal evidence against the null hypothesis, BF_01_ = 0.974.

In contrast, response times varied significantly as a function of condition, F(2,70) = 44.763, p < .001, η^2^_p_=0.561, BF_10_ = 1.659e+10. If only auditory information was available, participants showed the slowest response times (M = 814.08 ms, SD = 131.70). In line with a typical audiovisual facilitation effect, the fastest responses occurred if the information from both modalities was congruent (M = 700.78 ms, SD = 101.93), whereas response times for the trials with informative auditory but uninformative visual information (i.e., visually uninformative condition) ranged in-between (M = 742,01 ms, SD = 124.76). Pairwise-comparisons revealed that responses in the auditory-only condition were significantly slower compared to responses in the audiovisual congruent, Z = 5.20, *p*_corr_ < .001, r = 0.994, BF_10_=4987.07, and compared to responses in the visually-uninformative condition, t(35) = – 6.09, *p*_corr_ < .001, d = −1.02, BF_10_ = 28343.05. Responses in the audiovisual-congruent condition were significantly faster than responses in the visually-uninformative condition, *t*(35) = −4.39, *p*_corr_ < .001, d = −0.732, BF_10_ = 249.946.

To complement the analysis of response times and accuracy, we fit a drift diffusion model to the data (Voss et al., 2013). The pattern of drift rates, our primary modeling parameter of interest, nicely aligns with the above-reported pattern of response times such that higher drift rates (i.e., higher rates of information uptake) were present in conditions with faster response times. A rmANOVA revealed significant differences in drift rate between conditions, F(2,70) = 35.654, *p* < .001, η^2^_p_= 0.505, BF_10_ = 3.201e+8. That is, on average, drift rate was the highest if both the visual and the auditory modality provided informative input (audiovisual-congruent, M = 3.13, SD = 0.82), whereas it was significantly lower if only auditory information was provided (MDN = 2.18), *p*_corr_ < .001, r = −0.988, BF_10_ = 2188.37, or when information from the visual modality was uninformative (M = 2.76, SD = 0.89), *t*(35) = 3.88, *p*_corr_ < .001, d = 0.646, BF_10_ = 64.70. In addition, drift rates in the auditory-only condition were significantly lower compared to the visually-uninformative condition, *t*(35)= −4.768, *p*_corr_ < .001, d = −0.795, BF_10_ = 699.54.

In addition to drift rate, parameters recovered by the drift diffusion model were threshold separation (a: M = 1.78, SD = 0.71), non-decision time (t0: M = 0.43, SD = 0.08), and the variability of non-decision time (st_0_: M = 0.27, SD = 0.11). Those parameters were not allowed to vary between conditions (see 4.9.2).

### 2.2 Visual speech information does not modulate ERP correlates of auditory attentional orienting

To investigate the impact of visual speech information on auditory attentional orienting, we inspected the N2ac component as a correlate of the deployment of covert auditory spatial attention (Gamble & Luck, 2011; Klatt et al., 2018b). Figure 3A illustrates the contralateral versus ipsilateral waveforms at anterior (FC3/4, C3/4, C5/6, FT7/8) electrode sites for the three conditions. The waveforms start to diverge at around 200 ms, emerging into an N2ac component. The latter is followed by a sustained anterior contralateral negativity (in the following referred to as SACN). Notably, in contrast to other studies on attentional orienting in the visual domain (i.e, investigating the N2pc component -the visual analogue of the N2ac; see e.g., Jolicœur et al., 2008), the contralateral and ipsilateral curves do not converge, indicating an end of the N2ac, but rather seamlessly merge into a sustained contralateral negativity.

**Figure 2.**
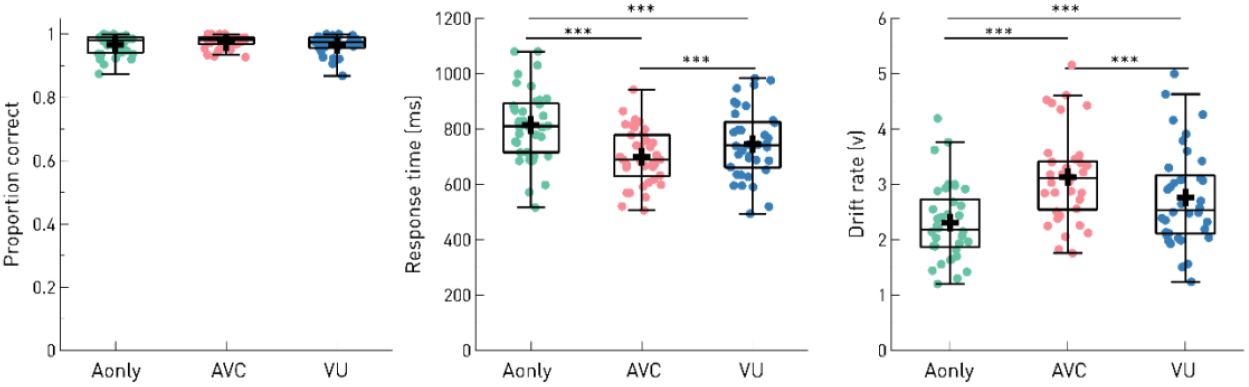
Behavioral performance. Proportion of correct responses (left panel), response times (middle panel), and drift rate (right panel) separately for each condition (Aonly = auditory only, AVC = audiovisual congruent, VU = visually-uninformative). Boxplots illustrate the interquartile range and the median. Whiskers extend to 1.5 times the interquartile range. The dots indicate individual participants’ mean scores per condition. A black cross denotes the condition mean across subjects.

**Figure 3.**
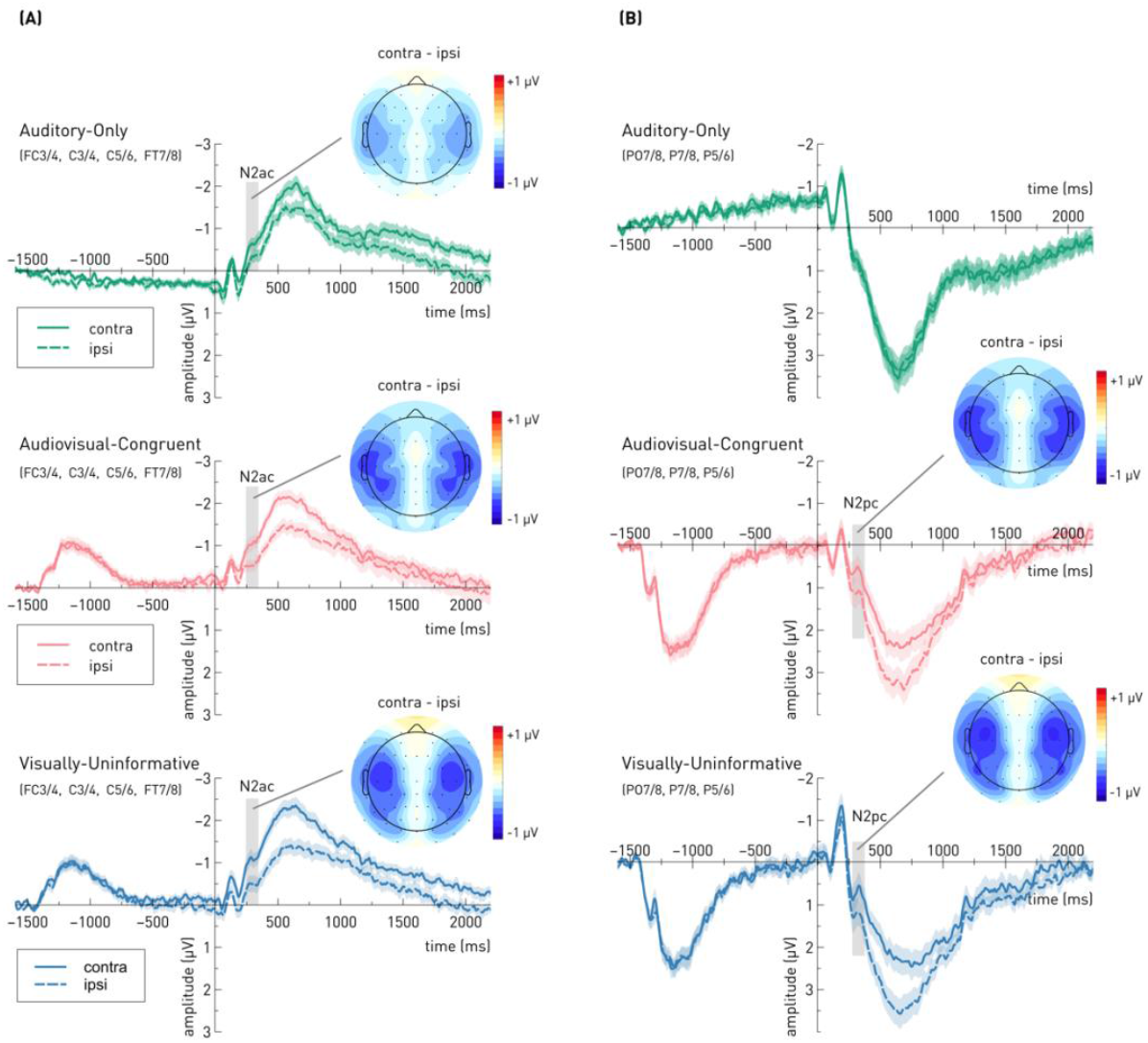
Event-related potential waveforms, contralateral and ipsilateral relative to the target sound and time-locked to sound-onset for each of the three experimental conditions. The left (A) and right (B) panel illustrate the ERPs for a cluster of anterior and posterior electrodes, respectively. Grey rectangles indicate the analysis time window. Scalp topographies show the distribution of voltages based on the contralateral - ipsilateral differences in the analysis time window. Consequently, the voltage is zero at the midline electrodes. The subtraction was mirrored across the midline to obtain a full topography; thus, the topographies are symmetrical.

Following up on the observed behavioral audiovisual facilitation effect, we hypothesized that if complementary visual cues facilitate target individuation and selection from a complex scene, this should in turn result in enhanced auditory attentional engagement. That is, we expected N2ac amplitudes to be greater in the audiovisual-congruent compared to the auditory-only condition. Further, a comparison of the auditory-only and the visually uninformative condition allows us to distinguish whether this multisensory facilitation results from the presence of visual input *per se* or whether it depends on the presence of meaningful (and thus, complementary) visual information.

Contrary to our expectations, a repeated measures ANOVA of mean N2ac amplitudes showed no significant differences in N2ac magnitude, χ^2^(2) = 1.50, *p* = .472, W = 0.021. A complementary Bayesian analysis of variance suggests that the data are approximately six times more likely under the null hypothesis than under the alternative hypothesis, BF_01_ =5.71. Yet, one sample *t*-tests against zero confirmed that an N2ac component reliably emerged in the audiovisual-congruent (MDN = −0.25), *p* = .005, *p*_corr_ = .014, r = −0.538, BF_10_ = 20.01, and the visually-uninformative condition (M = −0.59, SD = 0.82), *t*(35) = −4.29, *p* < .001, *p*_corr_ < .001, d = −0.714, BF_10_ = 189.30. In the auditory-only condition, the *t*-test against zero was only significant prior to correction for multiple comparisons, (MDN = −0.18), *p* = .033, *p*_corr_ = .060, r = −0.408, BF_10_ = 5.206; however, the Bayes factor provides moderate evidence in favor of the alternative hypothesis.

### 2.3 Audiovisual events automatically induce both auditory and visual selective attentional orienting

To investigate whether audiovisual events would automatically induce both auditory and visual attentional orienting, in addition to the N2ac, we examined the N2pc component at a cluster of posterior electrodes (PO7/8, P7/8, P5/6, see Figure 3B) as a correlate of visuo-spatial attentional selection (Eimer, 1996; Luck & Hillyard, 1994). While auditory attentional selection is required in any case given that the auditory input always provided task-relevant information, visual speech was not always informative. Hence, it could be that a-priori-knowledge about the lack of meaningful information in the visual domain results in the suppression of visual attentional spatial orientation, which should be indicated by the absence of an the N2pc component in the visually uninformative condition or an attenuation relative to audiovisual-congruent condition. However, based on a multisensory account of object-based selective attention, we expected that visuo-spatial attentional orienting should be evident, irrespective of the informational content of visual speech. That is, the N2pc component should not only be elicited in the audiovisual congruent, but also the visually uninformative condition. As can be seen in Figure 3B, this was in fact the case. One sample *t*-tests against zero verified that an N2pc reliably emerged in both the audiovisual-congruent (M = −0.51, SD = 0.99), t(35) = −3.113, *p* = .004, *p*_corr_ = .010, d = −0.52, BF_10_ = 10.01, and the visually-uninformative condition (MDN = −0.56), *p* < .001, *p*_corr_ = .007, r = −0.61, BF_10_ = 87.22. However, a paired sample *t*-test of mean N2pc amplitudes revealed no significant differences in N2pc magnitude between the audiovisual congruent and the audiovisual unspecific condition, t(35) = 0.27, *p* = .788, d = 0.05. A corresponding Bayesian analysis suggests that the data are approximately five times more likely under the null hypothesis than under the alternative hypothesis, BF_01_ = 5.40.

In the auditory-only condition, given that there was no visual information to attend to, we did not expect to see an N2pc component. However, one could argue that in natural conversations auditory speech typically goes along with a concurrent visual representation (i.e., when listening to a talker’s voice we can often look at their face and mouth movements). Hence, visually attending to a sound source location might be inherent to attentional orienting in complex scenes involving speech stimuli. To verify that the mere presence of a blank screen is not sufficient to elicit an N2pc component (see deviations from pre-registered methods, section 4.10), we conducted a one sample *t*-test for the auditory-only condition, revealing that the N2pc was – in fact – absent (M = −0.08, SD = 0.59), t(35) = −0.80, *p* = .428, *p*corr = .785, d = −0.134. Consistently, a complementary Bayesian analysis indicates that the data are approximately four times more likely under the null hypothesis than under the alternative hypothesis, BF_01_ = 4.14.

### 2.4 Temporally coherent visual cues do not speed up event-related lateralizations

A facilitation of audiovisual attentional orienting cannot only be evident in terms of a qualitative enhancement of attentional engagement (i.e., increased N2ac or N2pc amplitudes), but might also speed up attentional processing. This should be evident in earlier N2ac or N2pc onset latencies. Contrary to our analysis plan, onset latencies could not be separately assessed for the N2ac and the N2pc component using a fractional area latency (FAL) analysis, as the latter requires the component of interest to be clearly isolated from overlapping components (see *deviations from preregistered methods*, section 4.10e). Instead, Figure 3 illustrates, that the N2ac and the N2pc component seamlessly merged into a sustained contralateral negativity (SACN and SPCN, respectively). Thus, accounting for the overlap between the N2ac/N2pc component and the following sustained asymmetry (i.e., SACN / SPCN), the FAL was measured in-between 0 and 1100 ms relative to sound onset (i.e., including the post-stimulus interval up to the point at which on average a response was given in all conditions). Specifically, the points in time at which the area under the difference waveform reached 30% (onset-latency), 50% (midpoint-latency), and 80% (offset-latency) were determined in combination with a jackknifing approach (Kiesel et al., 2008, see the methods section for details). We expected that the behavioral benefit of presenting congruent audiovisual speech should also be evident in earlier onset- or midpoint latencies. There appears to be a trend towards later midpoint and offset latencies at anterior electrodes in the auditory-only condition (see Table 1), but the statistical analysis revealed no significant differences – neither, at anterior nor at posterior electrode sites (all *p* > .567).

**Table 1.** Onset (30%-FAL), midpoint (50%-FAL), and offset (80%-FAL) latency.

| Anterior cluster<br>(FC3/4, C3/4, C5/6, FT7/8) | Condition | 30%-FAL (ms) | 50%-FAL (ms) | 80%-FAL (ms) |
| --- | --- | --- | --- | --- |
|  | Aonly | 414 | 600 | 859 |
|  | AVC | 439 | 577 | 808 |
|  | VU | 422 | 569 | 807 |
| Posterior cluster<br>(PO7/8, P7/8, P5/8) | Aonly | - | - | - |
|  | AVC | 492 | 614 | 807 |
|  | VU | 489 | 614 | 812 |

*Note.* To obtain FAL, the area under the curve was measured in-between 0 and 1100 ms relative to sound onset. This includes the post-stimulus interval up to the point at which on average a response was given in all conditions. The table states the FAL-values obtained from the grand-average waveforms. In accordance with the pre-registration, no latency measures were extracted for the auditory-only condition at the posterior cluster, as we did not expect to find any posterior ERL in the absence of visual input. Aonly = auditory-only, AVC = audiovisual congruent, VU = visually-uninformative.

### 2.5 Exploratory ERP analysis reveals cross-modal interactions in working memory

As noted above, the N2pc was followed by a sustained posterior contralateral negativity (SPCN), while the N2ac was followed by a sustained anterior contralateral negativity (SACN). In previous visual search studies, the SPCN has been associated with subsequent in-depth analysis of the selected object in working memory (Jolicœur et al., 2008; Mazza et al., 2008; Töllner et al., 2013). Here, we show for the first time an analogous, lateralized auditory correlate and suggest that the SACN similarly reflects the engagement of auditory working memory as an intermediate processing buffer. Hence, to explore potential differences associated with processing beyond the initial attentional selection, reflected by the N2ac and the N2pc component, we performed an exploratory cluster-based permutation analysis contrasting ERL amplitudes in a broader time-window in-between 0 and 1100 ms. Please note, since no visually specific processing was expected in the auditory-only condition, at posterior electrodes, this comparison only included the two audiovisual conditions. Critically, as the same type of auditory information is presented in all conditions, any condition differences can only be due to differences in the informational content of visual speech (i.e., absent, congruent or uninformative). Figure 4 depicts the respective contralateral minus ipsilateral difference waveforms for each condition. In the stimulus-locked waveforms, the analysis revealed a small cluster in-between 500 and 560 ms at anterior electrode sites (*p* = .012), indicating a stronger contralateral negativity (SACN) for visually-uninformative compared to auditory-only trials. No additional clusters were identified in the stimulus-locked waveforms, neither at anterior not at posterior electrode sites.

**Figure 4.**
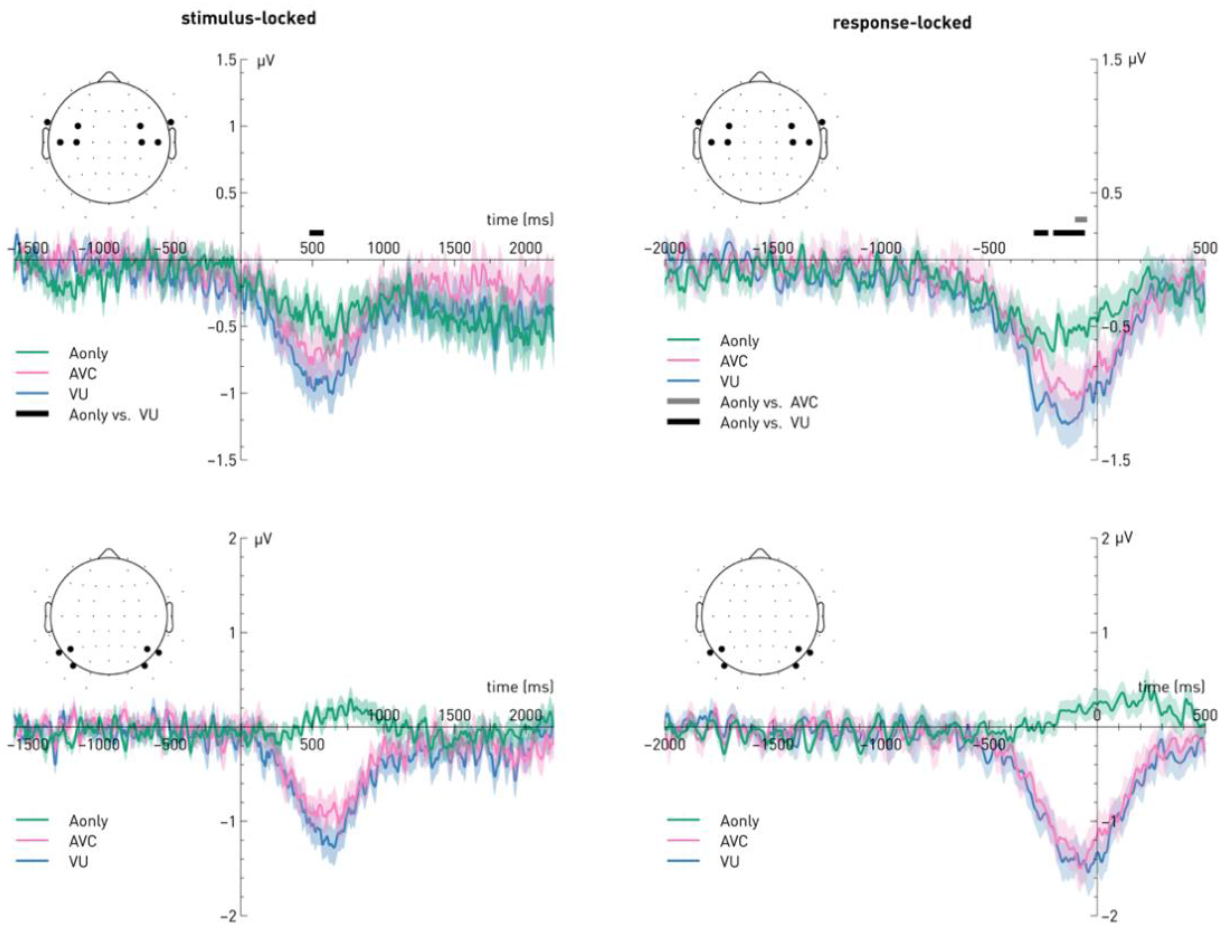
Event-related lateralizations (contra-ipsi) at anterior (top row) and posterior (bottom row) electrode sites. The left panel depicts the stimulus-locked waveforms relative to sound onset; the right panel depicts the response-locked waveforms time-locked to the response. Horizontal solid lines above the x-axis indicate the significant time points identified by a cluster-based permutation analysis. Aonly = Auditory-only, Avc = Audiovisual-congruent, Avu = visually-uninformative.

It is notable that, at anterior electrode sites, difference waveforms appeared to peak later in those conditions that also elicited slower response times, while the peak seemed to occur the earliest in the audiovisual-congruent condition which resulted in the fastest response times (see Table _1_ 1 and Figure 2). Consequently, we also inspected the response-locked waveforms. In fact, descriptively, the response-locked asymmetry was stronger in magnitude in all three conditions and both at anterior and posterior electrode sites. This suggests that both the SPCN and the SACN did vary with response-time, that is, the respective visual and auditory information is maintained in a spatially-specific fashion until the response was given. A cluster-based permutation analysis revealed three clusters at anterior electrode sites. Corroborating the stimulus-locked analysis, two clusters, ranging from −280 ms to −240 (*p* = .016) and from −190 to −70 ms (*p* = .002), signify a significantly stronger contralateral negativity in the visually-uninformative condition relative to the auditory-only condition. In addition, a small cluster ranging from −90 to −60 ms, indicates a significantly stronger contralateral negativity for audiovisual-congruent compared to visually unspecific trials (*p* = .024). The results suggest that dynamic audiovisual speech requires a greater cross-modal engagement of auditory working memory processes. Conversely, at posterior electrodes (i.e., related to visual working memory engagement), contrasting audiovisual-congruent versus visually-uninformative trials, no significant clusters were identified.

### 2.6 Time-frequency analyses reveal an impact on demands for cognitive control

Previous studies highlight the importance of synchronization of neural oscillations for multisensory processing (for a review, see Keil & Senkowski, 2018). Our own recent work, employing a similar audiovisual cocktail-party paradigm, suggests that in particular oscillatory power in the theta frequency over fronto-central scalp sites is modulated by the informational content of visual speech (Begau et al., 2022). Therefore, we performed an exploratory time-frequency analysis, using a cluster-based permutation analysis (see methods section for details). Figure 5 shows the time-frequency plots at a fronto-central electrode cluster for each condition as well as the differences between conditions for each of the three pairwise comparisons. In response to sound onset, an increase in theta power is visible in all conditions, followed by a broad suppression of power in the alpha and beta band. Notably, the increase in theta power is most pronounced in the auditory-only and the visually-uninformative condition, indicating increased demands for cognitive control, whereas it is substantially reduced when congruent information was presented in both modalities. The cluster-based permutation analysis confirmed this observation (see Figure 5D-E), identifying two significant clusters spanning the theta band (4-7 Hz) and slightly extending into the lower alpha range (up to 10 Hz) in the first ~330 ms (cluster 1, *p* < 10^-4^; see Figure 5D) and ~285 ms (cluster 5, *p* = 9.15^-4^, Figure 5F) following sound onset. Following the initial increase in power in the theta band, theta power drops in all three conditions; yet, while there is a steep decline in the auditory-only condition, reaching its minimum at around 500 ms post-sound onset (i.e., towards the end of the sound presentation), theta power bumps up again in the two audiovisual conditions at around 500 ms, indicating the need for sustained cognitive control when the audiovisual stimuli keep evolving across time. The latter observation is corroborated by the cluster-based permutation analysis, revealing two significant clusters; again, the clusters span the theta frequency range but slightly extend into the lower alpha band. When contrasting the auditory-only versus the audiovisual-congruent condition, the cluster ranges from ~300 to ~820 ms (*p* < 10^-4^), whereas for the contrast between the auditory-only versus the visually-uninformative condition, it ranges from ~350 to ~770 ms (*p* = 2.93^-4^). Finally, the contrast between the auditory-only and the audiovisual-congruent condition revealed a small but significant difference in a late cluster focused on theta frequency range, ranging from ~ 1030 ms to the end of the epoch at ~1780 ms (*p* = .004). For a close-up view of the temporal theta power dynamics and the topographic distribution of the described clusters, see Figure 6.

**Figure 5.**
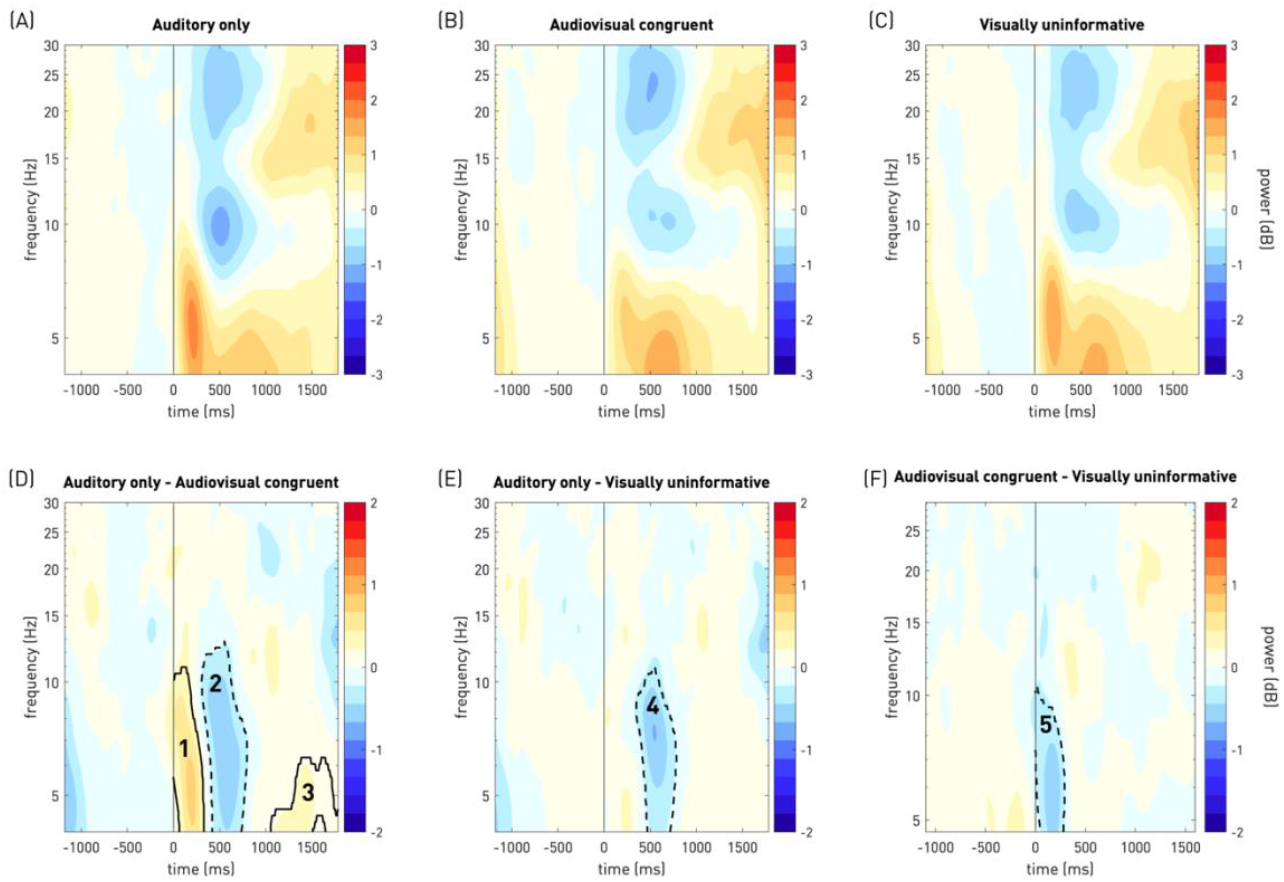
Time-frequency plots. The top row shows the time-frequency results separately for each condition. Power was averaged across a fronto-central electrode cluster, including electrodes Fz, Cz, FC1, FC2. The bottom row illustrates the differences between conditions. Solid lines indicate significant clusters with t-mass value greater than the 97.5th percentile of the null distribution, dashed lines indicate significant clusters with a t-mass value smaller than the 2.5^th^ percentile of the null distribution.

**Figure 6.**
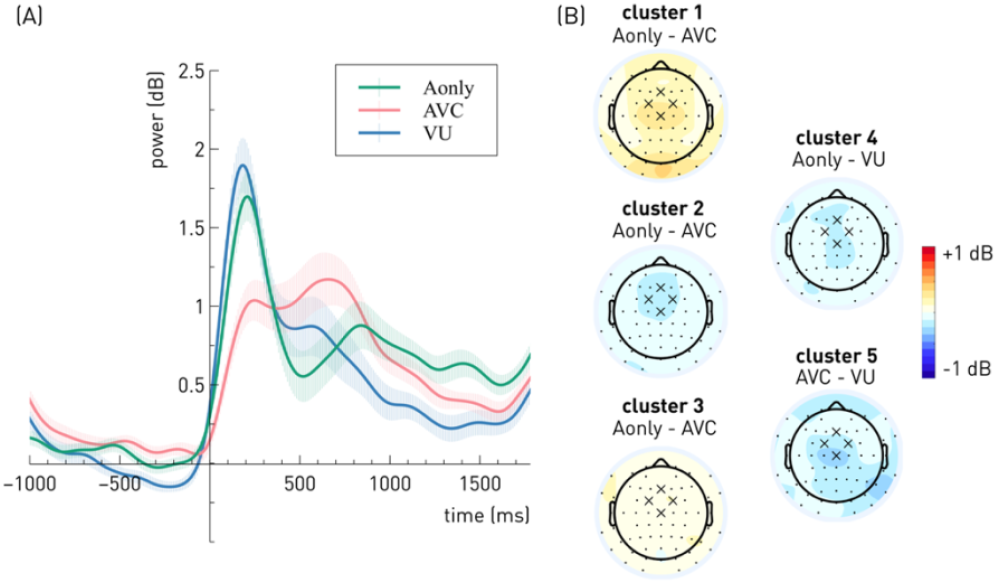
(A) Time-course of theta power (4-7 Hz) at frontocentral electrodes (FC1, FC2, Cz, Fz). (B) Scalp topographies associated with the clusters identified by the cluster-based permutation procedure, depicted in the same order as they appear in Figure 6 from left to right. Scalp topographies illustrate the mean difference in theta power (4-7 Hz) between conditions, averaged across the time window spanned by each cluster.

## 3. Discussion

When selecting task-relevant information from an audiovisual scene, theories of object-based attention predict that attention spreads across modalities to also encompass simultaneous signals from another modality, even when they are not task-relevant (Busse et al., 2005). Here, using ERP correlates of visual and auditory selective spatial attention, we investigate the interplay between modalities when selecting audio(-visual) speech information with a multi-talker scenario, while modulating the informational content of visual speech. In line with an object-based attention account, the central finding is that when confronted with an audiovisual scene, auditory attentional orienting was always accompanied by visual attentional orienting, irrespective of whether the visual modality provided task-relevant information or not. Critically, the lack of an N2pc component in the auditory-only condition suggests that the involvement of visual spatial attention is not *per se* elicited by any auditory stimulus, but depends on the availability or expectation of visual information that – in principle – forms a coherent percept with the incoming auditory information. That is, the mere presence of the blank screen as a potential ‘target’ for visual spatial attention in the auditory-only condition was not sufficient to elicit visual-spatial orienting. Beyond that, cross-modal interactions concerned measures of behavioral performance, working memory maintenance, and cognitive control.

### 3.1 Visual speech in the periphery facilitates sound localization within a multi-speaker mixture

Participants were able to identify and localize the target numeral most rapidly, when congruent audiovisual speech was present. Consistently, congruent audiovisual speech resulted in higher drift rates, indicating that the complementary visual information did in fact aid the evidence accumulation process, and suggesting that drift rate presents a sensitive and reliable measure of multisensory benefits (Chau et al., 2021; Murray et al., 2020). Overall, the behavioral results are consistent with the well-established audiovisual facilitation effect, demonstrating that bimodal stimulus presentation facilitates recognition accuracy, speeds up response times (Molholm et al. 2004, Giard & Peronnet, 1999, Miller, 1982), and improves perceptual sensitivity (Wahn et al. 2017). Converging effects have been reported specifically for speech comprehension, where the presence of congruent visual speech information improves comprehension compared to auditory-only speech (Bernstein et al., 2004; Tye-Murray et al., 2011). Notably, audiovisual facilitation and, more specifically the effects of visual speech on speech comprehension, have been predominantly investigated while presenting a single articulating speaker in central (i.e., foveal) vision. Hence the present findings extend this literature, illustrating that the positive effect of visual speech information on speech intelligibility and comprehension hold even when not directly gazing at a speaker’s mouth (for converging findings see Kim & Davis, 2011).

Interestingly, and contrary to our expectations, we found that uninformative visual speech still led to faster responses and higher drift rates compared to the auditory-only condition. Yet, the results are plausible, considering that visually unspecific speech – even though uninformative in terms of speech content – still conveyed temporal information that can increase the sensitivity to upcoming sensory information (Peelle & Sommers, 2015). Consistently, previous studies have shown that degraded visual speech (Tye-Murray et al., 2011), the presentation of synthetic visual stimuli like a dynamic mouth-like shape (Bernstein et al., 2004; Tye-Murray et al., 2011), or a dynamic rectangle whose horizontal extent was correlated with the speech envelope (Bernstein et al., 2004) still improved speech detection thresholds compared to auditory-only speech. More generally, this supports the idea that binding between features across modalities at a perceptual level (i.e., resulting in the percept of a multisensory object) relies most strongly on temporal coherence rather than on other consistencies, such as phoneme-viseme relationships (e.g., presentation of a /u/ vowel paired with an image of protruded lips; Bizley et al., 2016). Finally, it is noteworthy that the effect of uninformative visual speech on sound localization performance occurred even though we used a block-wise presentation of conditions, allowing participants to anticipate that visual input would remain task-irrelevant. As outlined above, this is in line with the concept of object-based attention.

### 3.2 Behavioral benefits do not rely on improved attentional selection

The behavioral benefits observed for congruent audiovisual speech raise the question to what extent visual cues aid the listener to individuate and select the task-relevant target from the mixture of speech streams. Therefore, we investigated the effect of congruent and uninformative visual speech information on N2ac amplitudes and latencies, a correlate of auditory attentional selection (Gamble & Luck, 2011; Klatt et al., 2018b). While an N2ac component emerged in all conditions, indicating that auditory spatial attention was focused on the location of the target numeral, N2ac amplitudes did – unexpectantly – not differ significantly between conditions; rather, a Bayesian analysis provided evidence in favor of the null hypothesis. Beyond that, the analysis of ERL latencies also failed to provide any evidence for differences in onset-, midpoint of offset latency between conditions. As a caveat, it should be noted though that the lack of a significant effect could not be substantiated by evidence in favor of the null hypothesis since it was unclear whether the jackknifing procedure is compatible with a Bayesian analysis of the data.

Overall, the present results suggest that auditory spatial attentional orienting was not influenced by the informational content of visual speech. Given that the type of auditory speech information was equivalent across all three conditions, always providing task-relevant input, this appears absolutely plausible. However, this does imply that the behavioral benefits emerging from temporally coherent visual cues do not rely on stronger attentional engagement. Although, it should be noted that attentional selection, as marked by the N2ac component, already requires the auditory (or audiovisual) speech streams to be segregated. Therefore, it remains possible that audiovisual speech, when bound into a multisensory object, is more easily segregable from competing stimuli (Lee et al., 2019) and thus, audiovisual facilitation might occur at an earlier level of processing.

### 3.3. Attentional orienting in dynamic audiovisual scenes is object-based

As noted above, in the two audiovisual conditions, a shift of auditory spatial attention was accompanied by a shift of visual spatial attention. This was marked by a pronounced sound-locked N2pc component. The presence of an N2pc component in the audiovisual-congruent *and* the visually-uninformative condition is in line with a multisensory account of object-based selective attention (Molholm et al., 2007), according to which attention not only spreads across different features within the same unimodal object, but also across sensory modalities. Critically, the magnitude of the N2pc component in the two audiovisual conditions was not modulated by the informational content of visual speech. That is, it occurred irrespectively of whether visual speech was congruent with auditory speech or unspecific. This null finding was underpinned by a Bayes Factor analysis, providing evidence in favor of the null hypothesis. Considering that the block-wise presentation of conditions allowed participants to know beforehand that the visual speech information did not convey any information regarding the target location, the lack of differences in N2pc amplitudes between the two audiovisual conditions was striking. Previously, it has been shown that stimulus processing can be gated within the cortex to suppress or facilitate information flow for a given modality (Foxe et al., 1998; Mazaheri et al., 2014); however, such tasks typically involve the presentation of unrelated, low-level stimuli (such as pure tones and orientations) and the explicit instruction to attend to only one modality. Here, ecologically valid stimulus material was used, showing a speaking face. In such natural conditions, it appears that visual attention is automatically deployed to the relevant target location, irrespective of whether the visual modality provides additional task-relevant information or not. This observation is in line with previous studies, illustrating that attention can spread across modalities to encompass task-irrelevant sensory features, even when presented at a discrepant location (Busse et al., 2005; Degerman et al., 2007; Van der Burg et al., 2011). For instance, Busse et al. (2005) asked participants to respond to visual targets in a stream of lateralized, flashing checkerboards, while half of the stimuli were paired with a centrally presented, task-irrelevant tone. The neural responses to the task-irrelevant tones were larger, when they co-occurred with an attended compared to an unattended visual stimulus. Specifically, this was evident in terms of enhanced auditory cortex activity and a sustained, enhanced ERP reflection over fronto-central scalp sites. The authors interpret their results as an object-based selection process, grouping the attended visual stimulus with the synchronous, initially unattended auditory stimulus. Analogously, in the present study, such automatic audiovisual grouping occurs for naturalistic audiovisual speech stimuli. Importantly, the present study used temporally coherent, ecologically valid recordings of the speakers’ faces, including auditory and visual speech information that actually originates from the same spatial source. Beyond that, the present study extends the findings by Busse et al. (2005), showing that principles of object-based attention hold when introducing competition between two concurrent talkers.

On a similar note, Van der Burg et al., (2011) demonstrated that a task-irrelevant auditory signal enhanced the neural response to a synchronized visual event. Notably, they showed that a sound-elicited N2pc component occurred in response to the audiovisual stimuli irrespective of whether the visual stimulus was a task-relevant target or a distractor. Although the latter study demonstrates that sound-induced visual attentional orienting can occur in an automatic fashion, it differs from the present study in that it uses a classical pip-and-pop effect paradigm (Van der Burg et al., 2008), presenting a single sound together with a dynamically changing visual search display, containing several competing visual items. In this setup, the N2pc effect is likely due to attentional capture given that the isolated auditory stimulus presents an odd-ball-like stimulus (see also Gao et al., 2021, argueing against a role of multisensory integration in the pip-and-pop paradigm). In contrast, in our paradigm, the two audiovisual speech streams are equally salient, the number of visual and auditory features is balanced, and thus, allows for the binding of audiovisual features into coherent and meaningful multisensory objects.

Along those lines, the fact that we observed an N2pc component for all audiovisual stimuli, irrespective of the informational content of visual speech, supports the notion that binding – that is, the formation of an audiovisual object representation – is a largely perceptual process rather than a cognitive process (Bizley et al., 2016). The latter is, as noted above, postulated to rely most strongly on temporal coherence rather than on higher-order consistencies (Bizley et al., 2016). The present results support this line of thought, for though subjects were previously informed that the lip movements would not match the presented auditory input in the visually-uninformative condition, attentional selection, when operating on the object-level, always incorporated both visual and auditory features.

A final, critical point concerns the lack of an N2pc component in the auditory-only condition, given that there was no visual information presented for subjects to attend to. What might seem trivial, is quite important, because it demonstrates that the involvement of visual spatial attention requires the presence or expectation of visual information that – in principle – forms a coherent percept with the incoming auditory information. That is, participants did not focus their visual attention on the blank screen. This is further in line with the notion that the N2pc reflects attentional deployment to objects rather than to spatial locations serving as placeholders for upcoming information (Woodman et al., 2009).

### 3.4 Congruent audiovisual information reduces demands for cognitive control

In line with the behavioral audiovisual facilitation effect, the observed theta power dynamics were indicative of reduced demands for cognitive control during the initial period of audiovisual processing. Specifically, following sound onset, the early increase in theta power was greatly diminished if both modalities provided congruent information, whereas a stronger increase in theta power was evident if acoustic speech was paired with visually uninformative input. This is consistent with previous work showing an association between activity in the theta-band and the processing of mismatching audiovisual speech stimuli (Keil et al., 2012) and the successful integration of audiovisual speech (Lindborg et al., 2019). Moreover, the finding add to a mounting body of literature, interpreting theta power as a mechanism for cognitive control (Cavanagh & Frank, 2014) and conflict processing (Morís Fernández et al., 2018; for a review see Keil & Senkowski, 2018). Notably, the present study also showed a stronger increase in central theta band power following auditory-only input compared to congruent audiovisual speech. Thus, it appears that the processing of auditory input posed similar demands for cognitive control as the incongruent (i.e., visually unspecific) condition. Although auditory-only speech did not require subjects to resolve a conflict arising between modalities, this is plausible because the task still required subjects to separate the relevant from the irrelevant speech stream; the response time (and drift rate) data clearly show that this was more challenging when only auditory information was available.

### 3.5 Audiovisual speech results in the enhanced cross-modal recruitment of auditory working memory

Target selection, as indicated by the N2ac and the N2pc component, was followed by a sustained contralateral negativity, reflecting the subsequent processing of selected objects in working memory (Berggren & Eimer, 2018). That is, the present work illustrates that just as visual search relies on working memory storage (Emrich et al., 2009; Luria & Vogel, 2011), searching for a relevant target within a multisensory scene recruits both visual and auditory working memory. Accordingly, the observed sustained asymmetry over posterior scalp sites corresponds to a SPCN (Jolicœur et al., 2008), which is typically found in the context of visual search and indexes post-selective recurrent processes that are recruited when the extraction of detailed or response-critical information from working memory is required (Schneider et al., 2014; Töllner et al., 2013). In addition, for the first time, we show an analogous sustained contralateral negativity over anterior scalp sites, which we refer to as SACN. The latter likely presents a lateralized signature of sustained processing in auditory working memory, analogous to the SPCN. Previous studies of auditory short-term memory maintenance have primarily used non-lateralized or bilateral sound presentation and reported a sustained anterior negativity at fronto-central scalp sites (Lefebvre et al., 2013; Nolden et al., 2013; for a review, see Nolden, 2015). Further studies will be required to investigate whether the lateralized SACN similarly increases with set size, reaching a plateau as working memory capacity is exhausted, as has been shown for established working memory measures (Alunni-Menichini et al., 2014; Luria et al., 2016; Vogel & Machizawa, 2004).

An exploratory analysis of ERL in a broad post-stimulus interval revealed that SACN amplitudes were larger when acoustic speech was paired with visually uninformative speech compared to auditory-only speech. Critically, since the same type of auditory information was presented in all three conditions, any modulations of SACN amplitude can only be due to changes in the visual input. Consequentially, we argue that the modulations of SACN amplitude clearly indicate cross-modal interactions in working memory. Larger SACN amplitudes in the visually uninformative condition are in line with previous observations in visual search experiments, showing that the magnitude of the sustained contralateral negativity depends on the degree of post-selective processing that is required to extract detailed object information from working memory representations (Luria & Vogel, 2011; Töllner et al., 2013). While the differences between conditions only occurred in a relatively small time-window in the stimulus-locked ERLs, a response-locked analysis showed that the effect strongly varied with response latencies. Accordingly, the overall amplitudes in the response-locked ERLs appeared to be larger. This observation is consistent with previous work in the visual domain, illustrating that the SPCN co-varies with response time (Schneider et al., 2014) and in a broader context, also corroborates the newly emerging perspective on working memory that emphasizes its role for guiding actions (Heuer et al., 2020; Rösner et al., 2022; Schneider et al., 2017; van Ede, 2020). The response-locked analysis further revealed an additional cluster related to larger SACN amplitudes in the audiovisual-congruent compared to the auditory-only condition shortly before response onset. Hence, overall, it appears that the additional presence of visual speech content results in a greater cross-modal engagement of auditory short-term memory processes than acoustic speech alone. This is likely – at least in part – due to the dynamic properties of the audiovisual speech material used in the present study. For instance, the lip-movement outlasted the auditory speech articulation (completed after on average 550 ms) by on average ~70 ms. Further, the fade-out of the video did not start until several hundred milliseconds later. Hence, as the (audio-) visual input continues to evolve across time, this poses prolonged requirements on working memory in order to maintain a coherent percept of the incoming speech signal. Consistent with this interpretation, the late oscillatory power dynamics in the theta band suggest that in particular the two audiovisual conditions require additional demands for cognitive control once the acoustic speech input fades away, but the visual stimuli are still evolving. Accordingly, the cluster-based permutation analysis revealed two late clusters (~300 – 800 ms), showing higher theta power for the audiovisual congruent as well as the visually-uninformative condition compared to auditory-only speech presentation.

### 3.6 Conclusion and future work

In summary, the present findings add to a growing body of research addressing the interplay between modalities when navigating in an essentially multisensory environment. We show that covertly attending to a task-relevant auditory speech stream was naturally accompanied by covert attention to the co-occurring visual speech input (i.e., the speaker’s face), even if the presented lip movements were essentially unrelated to the acoustic articulations, suggesting automatic audiovisual binding and object-based attentional orienting. Here, we argue that this effect relies on the largely preserved temporal coherence between visual and auditory speech in the visually uninformative condition. Future work could provide further insights regarding the audiovisual object properties required for spatial attention to spread across modalities by manipulating the temporal (a)synchrony between modalities and the perceptual complexity of the visual stimulus material.

Surprisingly, even though behavioral measures clearly indicated that both congruent and uninformative visual cues aided the listeners to localize a task-relevant sound, this behavioral audiovisual facilitation effect was not reflected in ERP correlates of attentional selection. Using dynamic recordings of naturalistic audiovisual speech stimuli, this extends the largely auditory literature on attentional orienting in complex auditory scenes and provides new insights on cross-modal interactions at the audiovisual cocktail-party.

## 4. Materials and methods

The hypotheses and methods were preregistered on the Open Science Framework (https://osf.io/vh38g) after data from six subjects had been collected, but before any of the behavioral or ERP data was observed or analyzed. Deviations from the pre-registered analysis pipeline are specified in the text and further explained in section 4.10. Additional exploratory analyses not included in the preregistration are described in section 4.11. The experimental procedure was conducted in accordance with the Declaration of Helsinki and approved by the Ethical Committee of the Leibniz Research Centre for Working Environment and Human Factors, Dortmund, Germany.

### 4.1 Power Analysis

The target sample size of 36 participants was derived while attempting to find a good balance between limited resources, theoretical justification, and optimizing power. Specifically, the sample size rationale focused on maximizing power to detect a potential latency effect, as we expected that out of all the dependent variables of interest, the latencies effects would be the smallest. In addition, we aimed for a target sample size divisible by 9 to be able to balance the target numeral across subjects.

First, we considered previous reports of N2pc latency effects in the literature (to our knowledge, there are no studies available that report N2ac latency effects). There is a substantial variance in the size of latency effects depending on the specific experimental manipulations, ranging from as little as 9 to up to 52 ms (Callahan-Flintoft & Wyble, 2017; Feldmann-Wüstefeld & Schubö, 2015; Foster et al., 2020; Kumar et al., 2009; Matusz & Eimer, 2013). Thus, considering that (1) N2pc latency effects in the published literature might be biased (i.e., over-estimates of the true effect size) and that (2) available studies are not necessarily similar enough to the present study, the expected effect size might be much smaller (e.g., ~10 ms, cf. Matusz & Eimer, 2013, Feldmann-Wüstefeld & Schubö, 2015). This is only 1/3 of the effect size that Kiesel et al. (2008) assumed as representative for their simulation study, which yielded a recommended sample size of 12 subjects (with 100 trials for each condition) to obtain adequate power for an N2pc latency effect. Consequentially, we aimed for a sample size of up to three times (i.e., 36) as large.

With this anchor point in mind, we used the Superpower package (v0.1.0; Lakens & Caldwell, 2021) in R (v3.6.1) to simulate power for small (~10 ms), intermediate (~20 ms) and large (~40 ms) latency effects (number of simulations = 1000, alpha level = .05). Data from a previously published auditory search task (Klatt et al., 2018a, cf. pre-cue, sound localization condition) provided estimates of the to-be-expected standard deviation and mean differences (30% FAL = 43 ms, SD = 65 ms; 50% FAL = 21 ms, SD = 69 ms). Assuming a relatively large mean latency difference (between the audiovisual congruent and auditory-only condition) of 40 ms, a standard deviation of 70 and a correlation of r = .5, a sample size of 36 subjects yields a satisfactory power of 95 % for a main effect of modality and ~81% power for the relevant follow-up pairwise comparisons, when considering corrections for multiple comparisons. A much more conservative scenario, assuming a very small latency difference of only 10 ms (see, e.g., N2pc literature above; keeping all other parameters constant), yielded insufficient power estimates (< 11%). Plotting a power curve revealed that such a scenario would require a sample size that is well beyond our available resources (> 250 subjects). Even if the standard deviation was assumed to be substantially smaller (i.e., 35) – for instance, because jackknifing and a high number of trials resulted in more accurate measurements – and the correlation between measurements was increased (r = .7), power for a significant main effect still remained low (58.8 %, for a sample size of n = 36). Finally, assuming an intermediate latency effect of 20 ms, a standard deviation of 35, and a correlation of r = .5, a sample size of n = 36 was estimated to result in 95% power for a main effect of modality, as well as ~81 % power for the relevant pairwise comparisons (again, considering corrections for multiple comparisons). Power estimates remained adequately high for a main effect of modality (i.e., at 87.4 %) if the standard deviation was increased to 40 but dropped below the desired power of at least 80% when assuming a standard deviation of 45 (keeping all other parameters constant). Taken together, a sample size of 36 was chosen, yielding adequate power for large and intermediate effects.

### 4.2 Participants

In total, 43 subjects participated in the experiment. 7 subjects were excluded due to technical problems with the experimental software. The final sample consisted of 36 subjects (18 female, 18 male). The mean age in the sample was 24.94 years (range: 19 – 35). 18 subjects reported to be right-handed, 8 to be left-handed. None of the participants reported any neurological or psychiatric illness. Prior to the beginning of the experimental procedure, participants were given written information about the study and provided written informed consent. At the end of the experiment, participants were compensated for their time in the laboratory with 10 Euros or one research credit (if requested) per hour.

### 4.3 Sensory and cognitive abilities

All participants completed a pure-tone-audiometry (Oscilla USB 330; Immedico, Lystrup, Denmark). The automated procedure included the presentation of eleven pure tones at varying frequencies (i.e., 125 Hz, 250 Hz, 500 Hz, 750 Hz, 1000 Hz, 1500 Hz, 2000 Hz, 3000 Hz, 4000 Hz, 6000 Hz, 8000 Hz). Hearing thresholds were ≤ 25 dB in the speech frequency range (< 4000 Hz) for all but one subject. The latter showed slightly increased hearing levels of 30 and 35 dB at 125 Hz and 750 Hz, respectively. Those deviations were considered negligible and the subject was included in the analysis.

To assess participants’ visual acuity, Landolt C optotypes were presented at a distance of 1.5 m. To obtain the average visual acuity, we took the common logarithm of the individual measures prior to averaging; subsequently, the antilogarithm of the resulting average value was computed (Bach & Kommerell, 1998). On average, visual acuity was 1.58 (SD = 0.11, range: 0.94 – 2.25).

Finally, to assess participants’ general ability to selectivity focus their attention, the auditory task in the subtest ‘divided attention’ of the TAP-M (test of attentional performance – mobility version 1.3.1) was conducted. The task requires participants to listen to a series of tones presented via headphones. The tones are presented for 433 ms with a SOA of 1000 ms. The tones rhythmically alternate between a low-pitch and a high-pitch tone. Subjects need to respond via button press to the rare occasions of the same tone being presented twice. Table 2 summarizes the relevant T-scores for commission errors, omission errors, and response times.

**Table 2.**
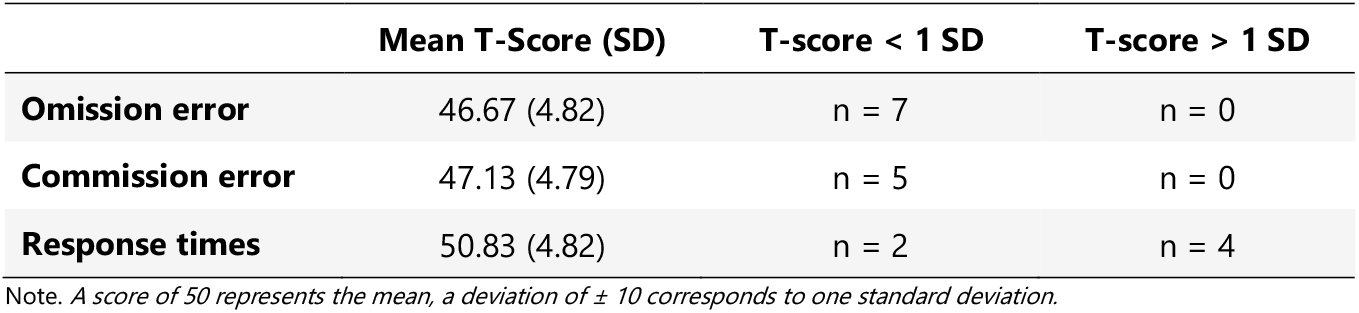
T-scores for the auditory task from the subtest divided attention of the TAP-M (v. 1.3.1).

|  | Mean T-Score (SD) | T-score < 1 SD | T-score > 1 SD |
| --- | --- | --- | --- |
| **Omission error** | 46.67 (4.82) | n = 7 | n = 0 |
| **Commission error** | 47.13 (4.79) | n = 5 | n = 0 |
| **Response times** | 50.83 (4.82) | n = 2 | n = 4 |

Note. A score of 50 represents the mean, a deviation of $\pm 10$ corresponds to one standard deviation.

### 4.4 Lipreading assessment

A short lip-reading assessment was conducted at the end of the experimental session. Participants watched muted video recordings of a single female speaker uttering five-word sentences (in German). All sentences were chosen from the *Oldenburger Satztest* (Wagener et al., 1999) and adhered to the following structure: Name, verb, number, adjective, object. Subjects were informed of the to-be expected structure of the sentences. In total, 30 sentences were presented. Prior to each sentence, two words were displayed (pre-cue). After each sentence, subjects were asked to indicate which one of the two pre-cue words the sentence contained (pre-cue task). Subsequently, a recognition probe was presented, and subjects were instructed to indicate whether the displayed word was present in the previous sentence or not. Oral responses were recorded by the instructor. See supplementary figure S1 for an illustration of the lipreading test procedure. For each task (pre-cue task vs. recognition task), a sum score was calculated, indicating the proportion of correct responses (i.e., ranging from 0 to 30). For one subject, the lipreading assessment could not be completed because of time constraints. In the pre-cue task, the average sum score was 23.63 (SD = 4.25, range: 15 – 30). In the recognition task, the average sum score was 17.37 (SD = 3.36, range: 11-25). Participants’ performance was similar to the performance of young adults in two previous studies, using the same test procedure (Begau et al. 2021, 2022).

### 4.5 Experimental paradigm

Participants performed a localization task. In each trial, participants were presented with two concurrent sound stimuli (recordings of German numerals ranging from 1 to 9) to the left and right of central fixation. The sounds were either presented (1) without any visual input (i.e., auditory-only), (2) in combination with a video of a speaker’s face and neck, providing congruent visual speech information (i.e., audiovisual-congruent), or (3) in combination with a video of a speaker, producing an unspecific “ba” lip movement; thus, the visual input did not provide any meaningful or task-relevant information (i.e., visually-uninformative). In all three conditions, participants were always required to indicate the location (left vs. right) of a prespecified target numeral by pressing the left or right button on a response pad, using the index and middle finger of their dominant hand. The target was presented on the left or right side in ⅓ of all trials (within a condition), respectively. In ⅓ of trials the target was absent. Target-absent trials did not require a button press. The pre-specified target numeral was balanced across subjects, such that each numeral served as the target four times.

The three conditions were presented in a block-wise fashion, with 288 trials each. The order of conditions was counterbalanced across subjects. In total, 864 trials were presented. Each trial had a total length of 2900 ms, followed by a 1000-ms inter-trial-interval. Figure 1B depicts an exemplary stimulus sequence. Please note that in each trial, two (audio-)visual stimuli were presented. In the two audiovisual conditions, the two monitors displayed a recording of a speakers face. Each trial started with 500 ms fade-in. Because the time between lip-movement onset and sound onset varied between the different speakers and stimuli, the fade-in was followed by a variable number of frames in which the speaker’s face remained still. This was done to ensure that sound onset was always at 1500 ms relative to trial onset. Following the articulation of the numerals, again, a variable number of frames, showing the speaker’s face with their mouth closed was shown to ensure an equal trial length. Each trial ended with a 400-ms fade-out. In auditory-only trials, the monitors remained black and displayed no visual input; otherwise, the trial timing remained unchanged.

### 4.6 Apparatus and stimulus material

The experiment is conducted in a dimly lit, sound attenuated room (5.0 × 3.3 × 2.4 m^3^). Pyramid-shaped foam panels on ceiling and walls as well as a woolen carpet on the floor result in a background noise level below 20 dB(A). Stimulus presentation was controlled using custom written Python software and three Raspberry Pi Modules (RPi 3 Model B) with a Raspbian GNU/Linux 10 OS, including the release Buster (Linux kernel version 5.4.83-v7+, processor: armv7l). Two 12” vertically aligned monitors (1080 x 1920 pixel resolution, 50 Hz refresh rate, Beetronics, Düsseldorf, Germany) were mounted to the right and left of central fixation on a horizontal array at a viewing distance of approximately 1.5 m, such that stimuli were presented at an eccentricity of approximately 3.4 ° visual angle (relative to central fixation). A full range loudspeaker (SC 55.9 – 8 Ohm; Visaton, Haan, Germany) was mounted underneath each monitor in a central position.

As stimulus material, a subset of video recordings from a previous study (Begau et al., 2021) were used. Specifically, the video recordings show the talker’s face and neck in front of a light-blue background (RGB 106, 145, 161) while uttering numerals, ranging from one to nine, or the word “ba”. The talkers were three native German females without any dialect and with an average fundamental frequency of 168 Hz (SE = 2.45), 218 Hz (SE =9.41), and 177 Hz (SE=16.09), respectively. All speakers served as the target speaker (i.e., speaker that utters the target numeral) equally often (i.e., in 1/3^rd^ of target-present trials). The same speaker never occurred in the same trial twice. That is, the speaker presenting the target numeral is paired with one of the two other speakers equally often. Critically, the occurrence of the three speakers, presenting the target numeral, was counterbalanced for a given target location (i.e., if the target occurs on the left side in 96 trials, each speaker serves as the ‘target speaker’ on 32 of those trials).

The video recordings were shot with 1920 x 1080-pixel resolution and a 50-fps frame rate under sound-attenuated conditions. Concurrently, mono audio tracks were recorded with a sampling rate of 48-kHz and 24-bit resolution, using a dynamic USB-microphone (Podcaster, RØDE, Silverwater, NSW, Australia). Audio tracks were noise reduced by 15 dB and normalized to −6 dB. Recording and editing of the audio tracks was done in Audacity(R) (version 2.3.0). To generate the stimulus material for the visually-uninformative condition, the video recording of the speakers uttering the word “ba” was paired with the audio tracks containing the numerals from one to nine.

Across the three speakers, the duration of the spoken numerals varied between 486 ms and 702 ms (M = 549.89 ms, SE = 8.33). The duration of visual speech, quantified as the time between the opening and closing of the talker’s mouth, ranged from 600 ms to 1520 ms (M = 1054 ms, SE = 43.12). On average, visual speech onset preceded auditory speech onset by 394 ms (range: 80 – 760 ms, SE = 33.93). For a detailed overview of the stimulus durations for each numeral and speaker, see supplementary table S1.

### 4.7 EEG data acquisition

The EEG signal was acquired from 64 Ag/AgCl electrodes (BrainCap, Brainvision, Gilching, Germany), distributed across the scalp according the extended internal 10-20 system. A NeurOne Tesla amplifier (Bittium Biosignals Ltd, Kuopio, Finland) was used to record the data at a sampling rate of 1000 Hz. AFz and FCz were used as online ground and reference electrode, respectively. During electrode preparation, the impedances were kept below 20 kΩ.

### 4.8 EEG preprocessing

EEG preprocessing was performed using MATLAB (R2021b) as well as the open-source toolbox EEGLAB (2021.1; Delorme & Makeig, 2004). The continuous EEG data was high-pass filtered at 0.01 Hz (pass-band edge; transition band width: 0.01 Hz, filter order: 330000, −6dB cutoff frequency: 0.005 Hz; see deviations from pre-registered methods, section 4.10a) and low-pass filtered at 40 Hz (pass-band edge; transition band width: 10 Hz, filter order: 330, – 6dB cutoff frequency: 45 Hz), using a using zero-phase, non-causal Hamming windowed sinc FIR filter. Bad channels were identified and rejected using the *pop_rejchan* routine implemented in EEGLAB. Specifically, channels with a normalized kurtosis greater than 5 standard deviations of the mean were excluded. To allow for a reliable estimation of eye-movement and blink related independent components (ICs), fronto-polar channels (FP1/2, F1/2, F3/4) were retained in the dataset and excluded from the automated channel rejection procedure. The number of rejected channels ranged from 0 to 7 (M = 4.92, SD = 1.69). Rejected channels were interpolated using spherical spline interpolation. Subsequently, the data was re-referenced to the average of all channels.

A rank-reduced independent component analysis (ICA) was applied for artefact correction. That is, to account for rank-deficiency that is introduced through interpolation of rejected channels as well as re-referencing to the average of all channels, the number of components to be decomposed was adjusted accordingly. ICA was run on the epoched data; epochs included latencies of −1600 up to 2200 ms relative to sound onset. To speed up ICA, it was applied on a subset of the data, downsampled to 200 Hz and including every second trial. Further, a 1-Hz high-pass filter was applied to the data prior to ICA, since it has been shown that high-pass filtering at ~1 Hz increases the percentage of ‘near-dipolar’ ICA components (Winkler et al., 2015). Finally, to further improve the signal-to-noise ratio in the dataset submitted to ICA, the automated trial rejection procedure *pop_autoręj* was applied (threshold: 1000 μV, probability threshold: 5 SD, maximum % of trials to reject per iteration: 5%). At this point, on average, 6.47 % (SD = 4.13) of trials were rejected. To identify ICs that are unlikely to reflect non-artefactual brain activity, the automated classifier algorithm ICLabel (Pion-Tonachini et al., 2019) was used. For each IC, ICLabel assigns a probability estimate to the classes ‘brain’, ‘eye’, ‘muscle’, ‘line noise’, ‘channel noise’, and ‘other’. ICs that receive a probability estimate of > 30% in the category eye or < 30% in the category brain were flagged for rejection.

The ICA decomposition and ICLabel estimates were copied to the dataset including all trials, with a sampling rate of 1000 Hz and high-pass filtered at 0.01 Hz. Then, epochs ranging from −1600 to 2200 ms relative to sound onset (see *deviations from pre-registered methods*, section 4.10c) were created and baseline-corrected using the first 100 ms of the epoch (i.e., – 1500 to −1600) as a baseline. Finally, ICs flagged for rejection were subtracted from the data. Following ICA-based artefact correction, trials with large voltage fluctuations (±150 μV) were removed (M = 2.66%, SD = 5.69; see deviations from pre-registered methods, section 4.10b). None of the trials had to be excluded due to premature responses (RT < 150 ms). After preprocessing, on average 89.86 trials (SD = 8.21), 92.89 trials (SD = 6.08), and 92.36 trials (SD = 6.06) per target location remained in the auditory-only, audiovisual-congruent, and the visually-uninformative condition, respectively.

### 4.9 Statistical analyses

Statistical analyses were performed in MATLAB (R2021b) and JASP (v0.16.3). The significance of all inferential tests was evaluated at an alpha level of 0.05. If the assumption of sphericity was violated (Mauchley’s test *p* < .05), Greenhouse-Geisser corrected *p*-values are reported. If the assumption of normality was violated (Shapiro-Wilk test *p* < .05), Friedman’s test and Wilcoxon signed rank test were conducted as non-parametric equivalents of repeated-measures analysis of variance (rmANOVA) and paired-sample *t*-tests. As effect sizes for rmANOVA and paired-sample *t*-tests, partial eta squared (η_p_^2^) and Cohen’s d are reported, respectively. For the calculation of Cohen’s d, the standard deviation of the difference scores was used as the denominator. In addition to inferential test statistics, the Bayes Factor is reported, if applicable, in order to quantify evidence in favor of the null hypothesis (see *deviations from preregistered methods*, section 4.10h). Complementary Bayesian analyses with a default Cauchy prior were conducted using JASP (v0.16.3). If applicable, post-hoc paired sample *t*-tests were corrected for multiple comparisons, using the Benjamini and Yekutieli (2001) procedure for controlling the false discovery rate (FDR). Corrected *p*-values are denoted as *p*_corr_.

#### 4.9.1 Behavioral task performance

To assess behavioral performance, mean response times, accuracy, and drift rate served as dependent variables. For model specifications of the drift diffusion model, see section 4.9.2. Accuracy measures, response times, and drift rate were submitted to a one-way repeated-measures ANOVA (or a non-parametric Friedman’s test, if applicable). For all analyses, stimulus modality served as the within-subject factor with three levels (auditory-only, audiovisual-congruent, Visually-uninformative). If applicable, follow-up paired sample *t*-tests (or a non-parametric Wilcoxon signed rank test) were conducted to resolve a significant main effect of stimulus modality. Respective *p*-values were corrected for multiple comparisons.

#### 4.9.2 Drift diffusion modeling

To further decompose the processes underlying the stream of information processing in the present audio(-visual) localization task, the diffusion model framework was applied. The diffusion model assumes that in a two-choice decision paradigm, information is continuously sampled until enough evidence for a particular decision has been accumulated. The slope of this diffusion process is termed drift rate and can be interpreted as the speed of information uptake (Voss, Voss & Lerche, 2015). The amount of evidence that is required for a response to be initiated is determined by the threshold separation a. The latter indicates the distance between the upper and the lower decision boundary. A potential bias towards one of the two decision boundaries is described by the starting point z. Non-decisional processes such as stimulus encoding and response execution are accounted for by estimating a response-time constant (t0).

Here, the diffusion model analysis was conducted using fast-dm-30 (Voss & Voss, 2007). To avoid contamination by outliers, the data were screened using a two-step procedure: First, trials with extremely fast (< 150 ms) and extremely slow (> 2000 ms) responses were excluded from the diffusion model analysis. Subsequently, based on the log-transformed and z-standardized data, trials with response times ±3 standard deviations of the mean (per participant and condition) were excluded. On average, 1.4 trials (SD = 1.5) per subject were excluded from the diffusion model analysis. The upper and lower decision boundary were assigned to correct and incorrect responses, respectively. Assuming an unbiased decision processes, starting point z was set to 0.5. The variability of the starting point (sz), variability of drift rate (sv), and differences in the speed of response execution (d), as well as the percentage of contaminants (p) were set to zero. While threshold separation (a), non-decision time (t_0_), and the variability of non-decision time(st_0_) were fixed across conditions, drift rate – as the primary parameter of interest – was allowed to vary across conditions (for a similar approach, see Experiment 1 in Ratcliff & McKoon, 2008). That is, in total 6 parameters were estimated. Parameter estimation was based on a maximum likelihood algorithm. That is, optimal parameters were obtained by maximizing the resulting log-likelihood value. An alternative model, allowing threshold separation, non-decision time, and variability of non-decision to vary between conditions, yielded inferior model fit (data not shown).

To assess model fit, deviations of the predicted values from the empirical values were graphically evaluated. Specifically, using the construct-samples routine implemented in fast-dm (Voss & Voss, 2007), 500 datasets per subject and condition were generated based on each individual’s parameter values and trial numbers. The 25%-, 50%-, and 75% quantile of the resulting distribution of stimulated response times were contrasted with the corresponding quantiles in the distribution of empirical response times. In addition, predicted accuracy (averaged across all 500 generated datasets per subject) was plotted against the observed accuracy values. The closer the data points are to the line of perfect correlation, the better the model fit. Data points in Figure 7 scatter close to the diagonal, indicating acceptable model fit.

**Figure 7.**
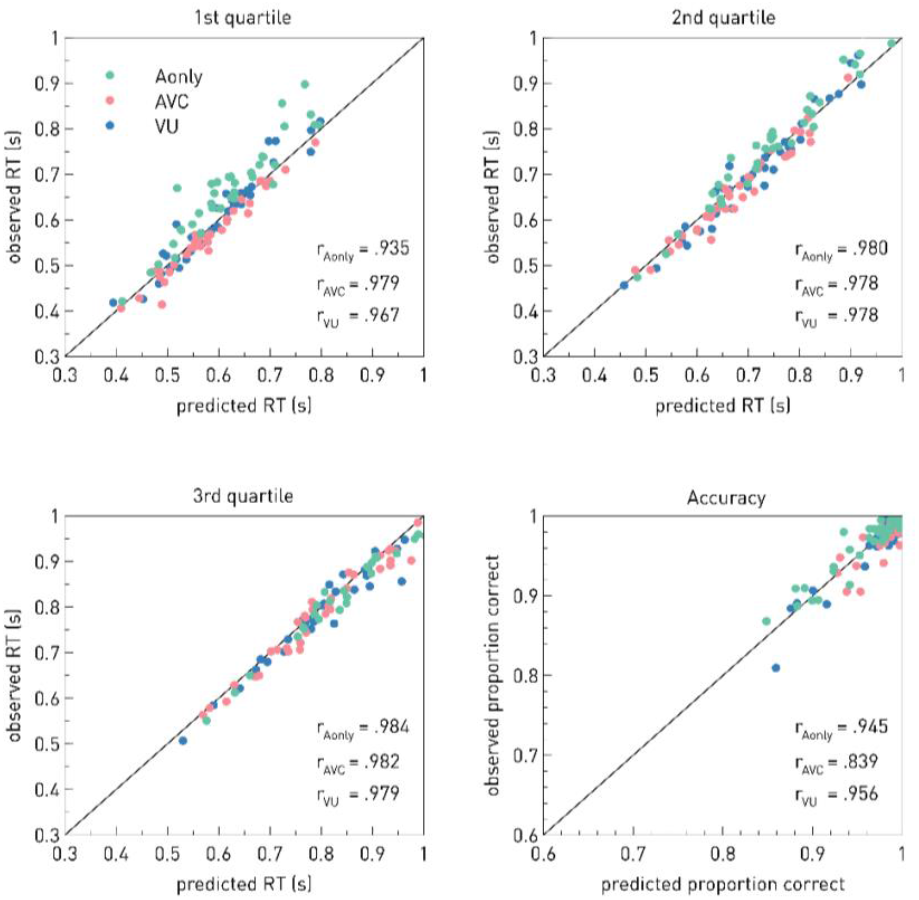
Graphical analysis of model fit. Scatter plots show the first three quartiles (.25, .5, .75) of the observed response time distribution as a function of the corresponding value from the predicted distribution as well as the observed proportion of correct responses as a function of the predicted proportion of correct responses. R denotes the corresponding Pearson correlation coefficient, separately for each condition (Aonly = auditory-only, AVC = audiovisual-congruent, VU = visually-uninformative).

#### 4.9.3 N2ac / N2pc amplitude effects

To determine the measurement time window for the quantification of N2ac and N2pc mean amplitude, we first determined the N2 peak latency in the grand average waveform (see supplementary figure S2) at an anterior (FC3/4, C3/4, C5/6, FT7/8) and posterior (PO7/8, P7/8, P5/6) cluster (see *deviations from preregistered methods*, section 4.10d). Subsequently, a 100-ms time window was set around the respective N2 peak. Accordingly, N2ac mean amplitude was measure in-between 244 to 344 ms; N2pc mean amplitude was measured in-between 275 to 375 ms. The electrodes of interest were derived from visual inspection of the scalp topographies, based on the grand-average contralateral minus ipsilateral difference waveforms (see supplementary figure S3). Note that we did not expect an N2pc to occur in auditory-only trials; thus, only audiovisual-congruent and visually-uninformative trials were collapsed for the inspection of the N2pc grand-average waveforms and topographies. In contrast, for the inspection of the grand average N2ac component, all three conditions were considered. N2ac amplitudes were submitted to a one-way repeated-measures ANOVA. Stimulus modality served as the within-subject factor with three levels (i.e., auditory-only vs. audiovisual-congruent vs. visually-uninformative). Follow-up paired sample *t*-tests were corrected for multiple comparisons. Finally, we tested for the general presence of an N2pc / N2ac effect within conditions (i.e., one sample *t*-test against zero).

#### 4.9.4 ERL latency effects

To quantitively assess the time-course of the event-related lateralization (ERL), the fractional area latency (FAL) technique was applied in combination with a jackknife approach (Kiesel et al., 2008). That is, first we created n average waveforms per condition, each of which was based on a subsample of n-1 (i.e., each participant was omitted from one of the subsample condition averages). Subsequently, for each of the subsample condition averages the point in time at which the negative area under contralateral minus ipsilateral difference waveform reaches 30% (i.e., 30%-FAL), 50% (i.e., 50%-FAL), or 80 % (i.e., 80%-FAL) of the total area (relative to a 0-mV baseline) was determined. The area under the curve was measured in-between 0 and 1100 ms, including the post-stimulus interval up until the point where on average the response had been given in all conditions. By including a larger time window than originally planned, we account for the fact that the N2ac/N2pc components overlapped with a sustained asymmetry (see *deviations from preregistered methods*, section 4.10e). The same electrode clusters as specified above were used. Onset latency measures from anterior electrodes sites (N2ac-SACN complex) were submitted to a repeated-measures ANOVA including the within-subject factor modality (i.e., auditory-only vs. audiovisual-congruent vs. visually-uninformative). For onset latencies from posterior electrode sites (N2pc-SPCN complex), a paired-sample *t*-test was conducted, contrasting the audiovisual-congruent and the visually-uninformative condition. *t* values and *F* values were corrected according to Kiesel et al. (2008; see also Miller et al., 1998). The corresponding *p*-values were obtained from the Student’s *t* cumulative distribution and the *F* cumulative distribution, respectively. As it is unclear to what extent jackknifing also inflates Bayes Factor estimates, we refrain from reporting the Bayes Factor for these analyses.

### 4.10 Deviations from preregistered methods

a. The preregistration stated that a high-pass filter of 0.1 Hz would be applied. The latter appeared to introduce a spread of evoked asymmetries in the post-stimulus interval into the pre-stimulus interval. Thus, the data was high-pass filtered at 0.01 Hz instead.
b. The preregistration stated that we would use the automated trial rejection procedure *pop_autoręj.* We did apply *pop_autoręj* on the subset of the data that underwent ICA decomposition. However, after ICA-based artefact correction the variance in the data is substantially reduced and consequentially, the iterative procedure that rejects data values outside a given standard deviation threshold results in the loss of large trial numbers, despite good data quality. Hence, following ICA-based artefact correction, only trials with large fluctuations (± 150 μV) were rejected using the algorithm *pop_eegthresh.*
c. Considering that the N2pc is a visually evoked potentials, we planned to conduct all N2pc analyses using ERP waveforms that were time-locked to lip-movement onset. However, the latter did not show any clear early evoked potentials (e.g., N1, P1). Thus, both N2pc and N2ac analyses were conducted, using the sound-onset locked waveforms. For transparency, the ERPs time-locked to lip-movement onset are displayed in the supplementary material.
d. The preregistration stated that we would measure mean N2ac and N2pc amplitude in a 100-ms measurement window centered around the 50% fractional area latency. The time window that serves as the boundary to measure the area under the curve was to be based on the visual inspection of the contralateral minus ipsilateral grand-average waveform as well in conjunction with considering typical time windows in the literature. Due to the overlap of the N2ac / N2pc effect with a late sustained asymmetry (see Figure), it was not possible to isolate the N2ac / N2pc and thus, applying the FAL technique to the data was not optimal. Instead, we chose to determine the N2 peak in the grand average waveform and then set a 100 ms time-window around the respective peak latency to compute mean amplitudes.
e. The preregistration stated that we would use the fractional area latency (FAL) technique to determine N2ac and N2pc onset latency. The time window that serves as the boundary to measure the area under the curve was to be based on the visual inspection of the contralateral minus ipsilateral grand-average waveform as well in conjunction with considering typical time windows in the literature. Due to the overlap of the N2ac / N2pc effect with a late sustained asymmetry (Figure 3), visual inspection of the grand average waveform does not allow us to isolate the N2ac / N2pc component and thus, determining an appropriate time window of interest to measure the area under the curve was difficult. Instead, we assessed the time-course of the entire event-related lateralization (ERL) in-between 0 and 1100 ms (i.e., including the post-stimulus interval up to the point where on average a response had been given in all conditions).
f. The preregistration stated that we would compute the 30% and 50% FAL. Considering that we are now measuring latency effects with respect to a much broader time window than anticipated, we decided to also compute the 80% FAL in order to adequately capture the entire time course of the N2ac-SACN and N2pc-SPCN complex, respectively, rather than just onset effects.
g. For the N2pc analysis, we initially planned to not consider the auditory-only condition at all because we did not expect an N2pc component to be present. However, to statistically verify that the N2pc component is in fact absent, we decided to conduct an additional one-sample *t*-test against zero for the N2pc amplitudes in the auditory-only condition.
h. In addition to the preregistered inferential statistics, we also report the Bayes Factor to be able to quantify evidence in favor of the null hypothesis.

### 4.11 Exploratory analyses

#### 4.11.1 Sustained ERL

To further explore the sustained asymmetry that overlaps with the N2ac and N2pc components, we performed a cluster-based permutation analysis to test for differences between the conditions. For the analysis of the stimulus-locked ERLs, we restricted the analysis on post-stimulus time-points up to the maximum average response time (i.e., 0 to 1100 ms, cf. Figure). In addition, we obtained response-locked waveforms from the data, ranging from −2000 ms to 500 ms relative to the response and performed the same analysis on the response-locked ERLs; only time points with 600 ms prior to the response were considered for the statistical analysis. Furthermore, we down-sampled the data to 100 Hz to reduce the total number of time points included in the analysis (Luck, 2014).

Specifically, for each pairwise comparison, the following steps were applied: First, at each time point a two-sided paired-sample *t*-test was conducted. The critical *t*-value to assess the significance at the cluster formation stage was 2.44. This corresponds to a *p*-value of .01 and sets a threshold for considering a sample point as a candidate member of a cluster. Using the Matlab function *bwconncomp(*), clusters in the original data were identified. For each cluster, the sum of all *t*-values was computed, constituting the cluster-*t*-mass. Then, to obtain a critical cut-off cluster-*t*-mass value that is required for a cluster in the data to be considered significant, we determine a null distribution from the observed data. That is, for each subject, the data were randomly assigned to the two conditions. Again, for the shuffled data, a paired sample *t*-test was conducted at each time point. This procedure was repeated a total of 10.000 times. On each iteration, samples with a *t*-value smaller than −2.44 or greater than +2.44 were selected and submitted to the function *bwconncomp().* For each negative (individual *t*-values < 2.44) or positive (individual *t*-values > 2.44) cluster, the sum of all *t-* values was computed. If several clusters were found, the largest cluster-*t*-mass value was stored. As a result, we obtained an estimate of the null distribution of maximum cluster t-mass values. At the inference stage, any cluster in the observed data with a cluster-*t*-mass value that is greater than the 97.5^th^ percentile of the null distribution, or smaller than the 2.5^th^ percentile of the null distribution was considered significant. This ensures an overall alpha level of 5%. The corresponding *p*-value was computed based on the inverse percentile of the observed cluster-level t-mass within the distribution of maximum or minimum cluster-t-mass values. More specifically, the *p*-value was calculated as (*inverse percentile /100*) or (*1 - inverse percentile/ 100*) for positive and negative clusters, respectively. If the observed cluster-level t-mass value exceeds the maximum or minimum cluster-level *t*-mass of the null distribution, the *p*-value is denoted as *p* < 10^-4^ (i.e., 1/number of permutation).

#### 4.11.2 Time-frequency analyses

To obtain event-related spectral perturbations (ERSPs, Makeig et al., 2004), we applied Morlet wavelet convolution. That is, the segmented EEG data was convolved with a series of complex Morlet wavelets that ranged in frequency from 4 to 30 Hz (increasing logarithmically in 52 steps). When creating the Morlet wavelets, the number of cycles is the crucial parameter that determines the width of the Gaussian that tapers the complex sine wave and thus determines the trade-off between temporal precision and frequency precision. Here, to obtain a good balance between temporal and frequency precision, the number of cycles in each wavelet was increased logarithmically as a function of frequency. The number of cycles at the lowest frequency was 3, the number of cycles at the highest frequency was 11.25. According to Grandchamp and Delorme (2011), a full-epoch length single-trial correction (i.e., using the full-trial length as a “baseline” period) was applied, prior to computing a common trial average pre-stimulus baseline. The time points in-between −500 to −100 ms relative to sound onset served as spectral baseline period.

Given that the time-frequency analysis was exploratory in nature, we performed a data-driven, non-parametric cluster-based permutation approach to test for differences between the condition. To increase the power to detect a true effect, we constrained the electrode sites and time points included in the analysis. That is, based on our previous study on audiovisual speech processing using similar conditions and stimulus material (Begau et al. 2022), we averaged the data across a cluster of fronto-central channels (including Cz, Fz, C1, C2). In addition, as we were not interested in anticipatory effects, only post-stimulus time points (i.e., in-between 0 and 1782 ms) were considered for the statistical analysis. That is, the data submitted to the cluster-based permutation test had the dimensions 52 (frequencies) x 182 (time points) x 36 (subjects). The same procedure as described in section 4.11.1 was followed.

## 5. Acknowledgements

This work was supported by a grant from the Deutsche Forschungsgemeinschaft (GE 1920/4-1).

## Supplementary material

### S1. Lip reading test procedure

**Figure S1.**
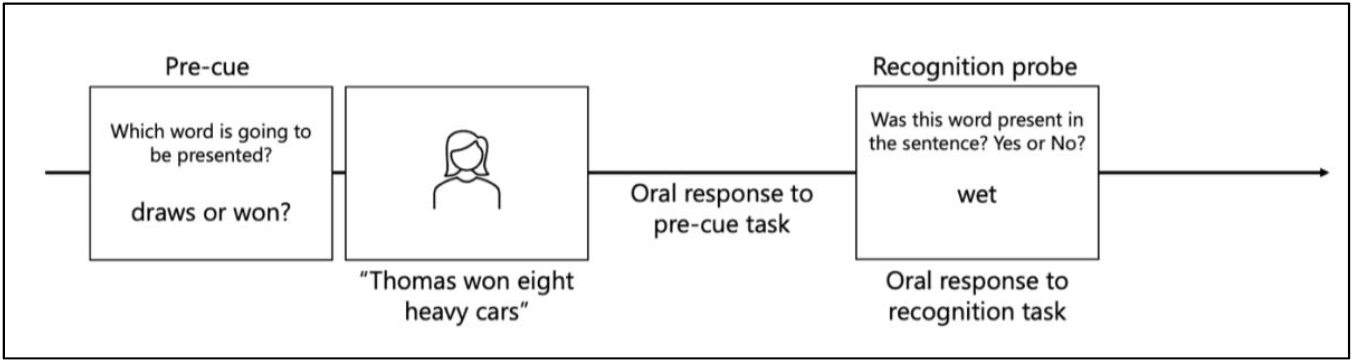
Lip reading test procedure. For illustrative purposes, the figure includes English sentences. However, the test was conducted in German. Following the pre-cue, participants were presented with a muted recording of a speakers face articulated a five-word sentence. All sentences had the same structure: Name, verb, number, adjective, object. Subsequently, subjects orally responded to the pre-cue task, indicating which of the two words they had previously seen (pre-cue) was included in the sentence. Finally, a recognition probe was presented. Participants responded by saying ‘yes’ or ‘no’. Oral responses were recorded by the experimenter.

### S2. Grand average ERP waveforms at anterior and posterior electrode clusters

Figure S2 depicts the grand average ERP waveforms at anterior and posterior electrode clusters that were used to derive the time window for statistical analyses of N2ac and N2pc mean amplitudes.

**Figure S2.**
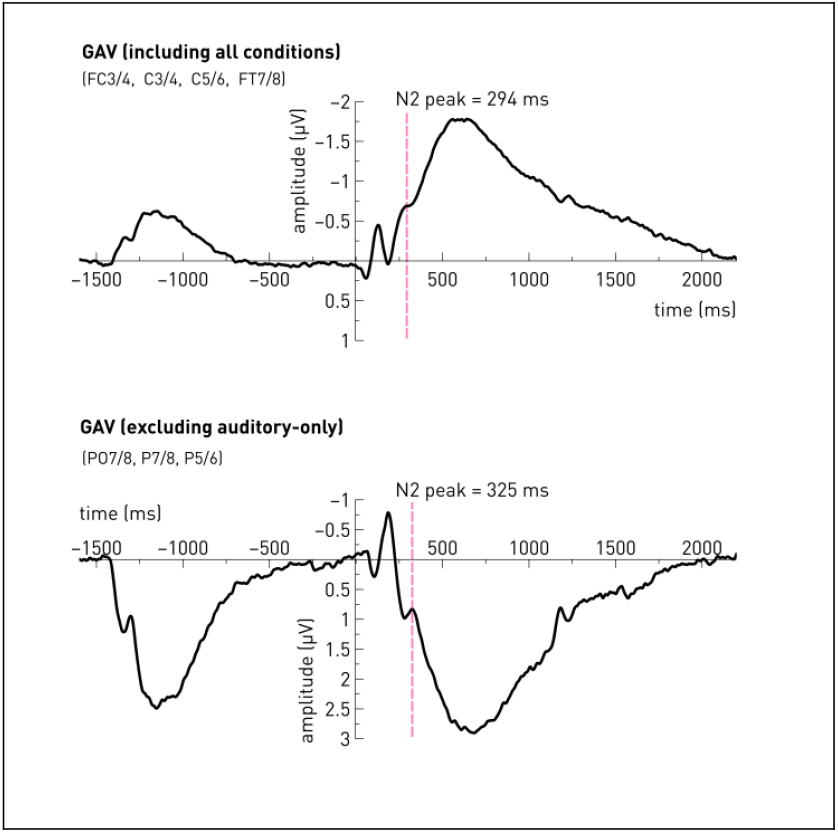
Grand average ERP waveforms at anterior (top) and posterior (bottom) electrode clusters. The time window for statistical analyses of N2ac and N2pc mean amplitudes was determined based off the N2-peak-latency in the grand average waveforms. That is, a 100 ms time window was set around the respective N2 peak latency. According to the preregistered procedure, for the inspection of N2pc waveforms, only the audiovisual congruent and the audiovisual-unspecific condition were included in the grand average. For the N2ac component, all conditions were averaged. The dashed line marks the N2 peak latency.

### S3. Grand average contralateral-ipsilateral scalp topographies

As indicated in section 2.8.3, the electrodes of interest were be derived from visual inspection of the scalp topographies, based on the grand-average contralateral minus ipsilateral difference waveforms. Supplementary figure S3 indicates the respective scalp topographies for a series of time windows. The top row includes the average across all three conditions (used to derive the anterior electrode cluster), the bottom row includes the average across the two audiovisual conditions (used to derive the posterior electrode cluster).

**Figure S3.**
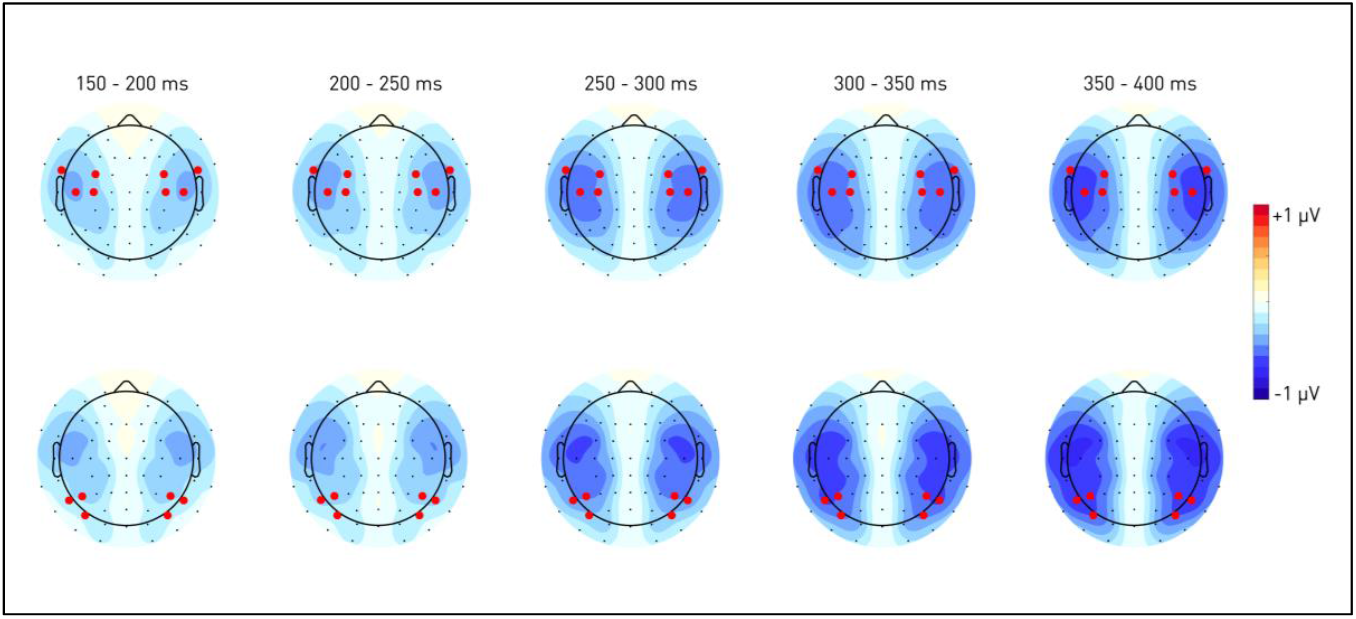
Time-series of scalp topographies. The chosen electrodes are highlighted in red.

**Table S1.** Stimulus properties.

|  | **Talker 1** |  |  | **Talker 2** |  |  | **Talker 3** |  |  |
| --- | --- | --- | --- | --- | --- | --- | --- | --- | --- |
| **Utterance** | Onset delay<br>lip-auditory | Length<br>auditory<br>stimulus | Length<br>visible lip<br>movement | Onset delay<br>lip-auditory | Length<br>auditory<br>stimulus | Length<br>visible lip<br>movement | Onset delay<br>lip-auditory | Length<br>auditory<br>stimulus | Length<br>visible lip<br>movement |
| **/Eins/** | 380 | 520 | 960 | 400 | 529 | 1000 | 760 | 595 | 1440 |
| **/Zwei/** | 580 | 532 | 1200 | 180 | 537 | 920 | 300 | 552 | 840 |
| **/Drei/** | 620 | 522 | 1260 | 500 | 504 | 1200 | 340 | 499 | 1020 |
| **/Vier/** | 280 | 551 | 900 | 560 | 607 | 1360 | 80 | 492 | 640 |
| **/Fünf/** | 220 | 575 | 860 | 580 | 702 | 1440 | 340 | 545 | 960 |
| **/Sechs/** | 300 | 569 | 1060 | 240 | 581 | 1020 | 560 | 584 | 1340 |
| **/Sieben/** | 200 | 525 | 740 | 280 | 566 | 880 | 540 | 546 | 1080 |
| **/Acht/** | 300 | 486 | 1040 | 760 | 563 | 1520 | 380 | 521 | 1100 |
| **/Neun/** | 120 | 587 | 820 | 580 | 532 | 1240 | 400 | 525 | 1000 |
| **Visually**<br>**unspecific**<br>**/Ba/** | 160 | see above | 600 | 660 | see above | 1360 | 220 | see above | 820 |

*Note.* The German utterances “Eins”, “Zwei” etc. as listed in the first column correspond to numerals one to nine. Denoted values are in ms. In the audiovisual-congruent condition, participants saw and heard matching visual and auditory speech utterances. In the audiovisual-unspecific condition, the auditory utterances were paired with a recording of the respective speaker’s face while saying “ba”. The same visual utterance was used for all nine visually unspecific stimuli per speaker. The onset of auditory speech was locked to always occur 1500 ms (i.e., 75 frames) after the beginning of the video. 1 frame corresponds to 20 ms (50 frames per seconds). The onset delay between the onset of the lip movement and the onset of auditory speech was computed determining the visual lip movement onset frame and subtracting it from 75. The length of the visible lip movement was computed by subtracting the visual speech onset and offset frame. Auditory speech duration was manually computed by selecting the visible portion of the acoustic sound wave in an audio editing software (Audacity v2.3.0).

